# Functional characterization of eicosanoid signaling in *Drosophila* development

**DOI:** 10.1101/2025.01.13.632770

**Authors:** Daiki Fujinaga, Cebrina Nolan, Naoki Yamanaka

## Abstract

20-carbon fatty acid-derived eicosanoids are versatile signaling oxylipins in mammals. In particular, a group of eicosanoids termed prostanoids are involved in multiple physiological processes, such as reproduction and immune responses. Although some eicosanoids such as prostaglandin E2 (PGE2) have been detected in some insect species, molecular mechanisms of eicosanoid synthesis and signal transduction in insects have not been thoroughly investigated. Our phylogenetic analysis indicated that, in clear contrast to the presence of numerous receptors for oxylipins and other lipid mediators in humans, the *Drosophila* genome only possesses a single ortholog of such receptors, which is homologous to human prostanoid receptors. This G protein-coupled receptor, named Prostaglandin Receptor or PGR, is activated by PGE2 and its isomer PGD2 in *Drosophila* S2 cells. *PGR* mutant flies die as pharate adults with insufficient tracheal development, which can be rescued by supplying high oxygen. Consistent with this, through a comprehensive mutagenesis approach, we identified a *Drosophila* PGE synthase whose mutants show similar pharate adult lethality with hypoxia responses. *Drosophila* thus has a highly simplified eicosanoid signaling pathway as compared to humans, and it may provide an ideal model system for investigating evolutionarily conserved aspects of eicosanoid signaling.

**Author Summary:** There are numerous bioactive lipids that control animal physiology, and some of them are commonly observed in both humans and insects. Well-studied insects such as fruit flies can therefore be an excellent gateway to learn about lipid molecules that have common functions in various animal species. In this study, we analyzed the fruit fly genome to look for genes that encode sensors (or receptors) for such lipid molecules called lipid mediators and found that the fruit fly only has one such receptor, as compared to ∼50 receptors in humans. Interestingly, the fly receptor we found was similar to human receptors for lipid molecules called prostanoids. Our work on the fruit fly prostanoid receptor further revealed that it is important for development of the fly respiratory system, and we showed that flies can also synthesize prostanoids in their body just like humans. Prostanoids are clinically important due to their pro-inflammatory functions, and some widely used drugs such as ibuprofen target enzymes that synthesize prostanoids. Our study on fruit fly prostanoids therefore provides a solid basis for using this simple organism to reveal common prostanoid functions in animals, which may provide important insights into animal health in general.

## Introduction

Eicosanoids are 20-carbon fatty acid-derived bioactive lipids that regulate multiple biological processes such as inflammation and reproduction in animals [1,2]. Eicosanoids are synthesized in response to both external and internal stimuli including injury, infection, and gestation cycle, and induce various biological responses [3]. For example, a subclass of eicosanoids termed prostanoids (prostaglandins, thromboxanes, and prostacyclins) are known as strong inflammation-inducing factors in mammals [4,5]. Prostanoids are synthesized from arachidonic acid (AA), a 20-carbon polyunsaturated fatty acid, through reactions mediated by multiple biosynthetic enzymes [2,6] and exert their biological functions by interacting with G protein-coupled receptors (GPCRs) [4]. Because of the pro-inflammatory functions of prostanoids, prostanoid biosynthetic enzymes are important targets of a class of widely used drugs known as non-steroidal anti-inflammatory drugs or NSAIDs [7,8].

In insects, several prostanoids, mainly prostaglandins, have been reported in some species [9,10]. Through applications of synthetic ligands and biosynthesis inhibitors, functions of prostanoids in insect immune responses, hemocyte migration, and ovarian development have been reported [11–13]. Recently, a GPCR has been identified as a prostaglandin E2 (PGE2) receptor that mediates its immunomodulatory activities in a few lepidopteran species [14,15]. Targeted knockdown of putative prostanoid synthases has also been shown to reduce prostanoid production as well as immune responses in a lepidopteran insect [16,17], providing first examples of genetic loss-of-function studies of eicosanoid signaling in insects.

In the fruit fly *Drosophila melanogaster*, genes that are potentially involved in prostaglandin signaling have been mostly studied in the context of follicle maturation and border cell migration in the ovaries [11,18,19]. However, to the best of our knowledge, prostaglandin molecules have rarely been reported in *D. melanogaster* to date, except a few studies that detected PGE2 immunoreactivity in adult fly homogenates [20,21] and various eicosanoid metabolites in the adult hemolymph [22]. Some studies even argue that *D. melanogaster* is unable to synthesize C20 oxylipins [23–25], and a C18 oxylipin-mediated inflammatory response pathway has been proposed [25]. Combined with the lack of information regarding any oxylipin receptors in *D. melanogaster*, our knowledge on eicosanoid signaling pathways in this important model organism is critically limited.

In the present study, we identified a single eicosanoid receptor-encoding gene in the *Drosophila* genome and investigated its functions during development. This GPCR, named Prostaglandin Receptor (PGR), can be activated by prostanoids *in vitro*, consistent with the fact that it is orthologous to mammalian prostanoid receptors. Detailed loss-of-function analyses and rescue experiments revealed that PGR is required for development of the adult tracheal system during metamorphosis. Moreover, through a comprehensive mutagenesis approach, we identified a PGE synthase that functions in the trachea and mediates local prostaglandin signaling that promotes adult tracheogenesis. Our study thus sets a solid basis and provides essential genetic tools for studying this highly conserved lipid signaling pathway in an important model organism.

## Results

### CG7497/PGR is a single prostanoid receptor in *D. melanogaster*

We first compared amino acid sequences of all Class A GPCRs in *D. melanogaster* and *Homo sapiens* to identify candidate receptors for lipid mediators in *D. melanogaster* (Fig 1A, S1 Table). Most fly receptors were clustered in accordance with their cognate ligands, and a single *Drosophila* receptor, CG7497, was clustered with *H. sapiens* prostanoid receptors. CG7497 orthologs are also conserved in other insect species (S1 Fig, S2 Table), and its lepidopteran orthologs have recently been reported as insect PGE2 receptors [14,15]. In contrast, no other *Drosophila* GPCRs were found orthologous to other human lipid mediator receptors, such as leukotriene B4 receptors and lysophosphatidic acid receptors (Fig 1A and 1B). This result suggests that insects have highly simplified eicosanoid signaling mediated by a single GPCR. In order to determine ligands for CG7497, the aequorin-based calcium mobilization assay was performed using *Drosophila* S2 cells. Aequorin, a calcium-dependent luminescent protein, and Gα15, a promiscuous G protein, were transfected into S2 cells along with a GPCR-containing vector (either CG7497 or *H. sapiens* PGE2 receptor HsEP2) or an empty vector (Mock). By quantifying calcium-dependent aequorin luminescence as a readout of Gα15-coupled GPCR activation [26], it was confirmed that CG7497 was strongly activated by PGE2 or PGD2, while it was only slightly activated by PGF2α, U46619 (thromboxane A2 analog), or iloprost (prostacyclin analog) [27,28] (Fig 1C). These results indicate that CG7497 is a single *Drosophila* prostanoid receptor that can be activated by a wide variety of prostanoids, particularly PGD2 and PGE2. Hereafter, we call CG7497 PGR.

**Fig 1.**
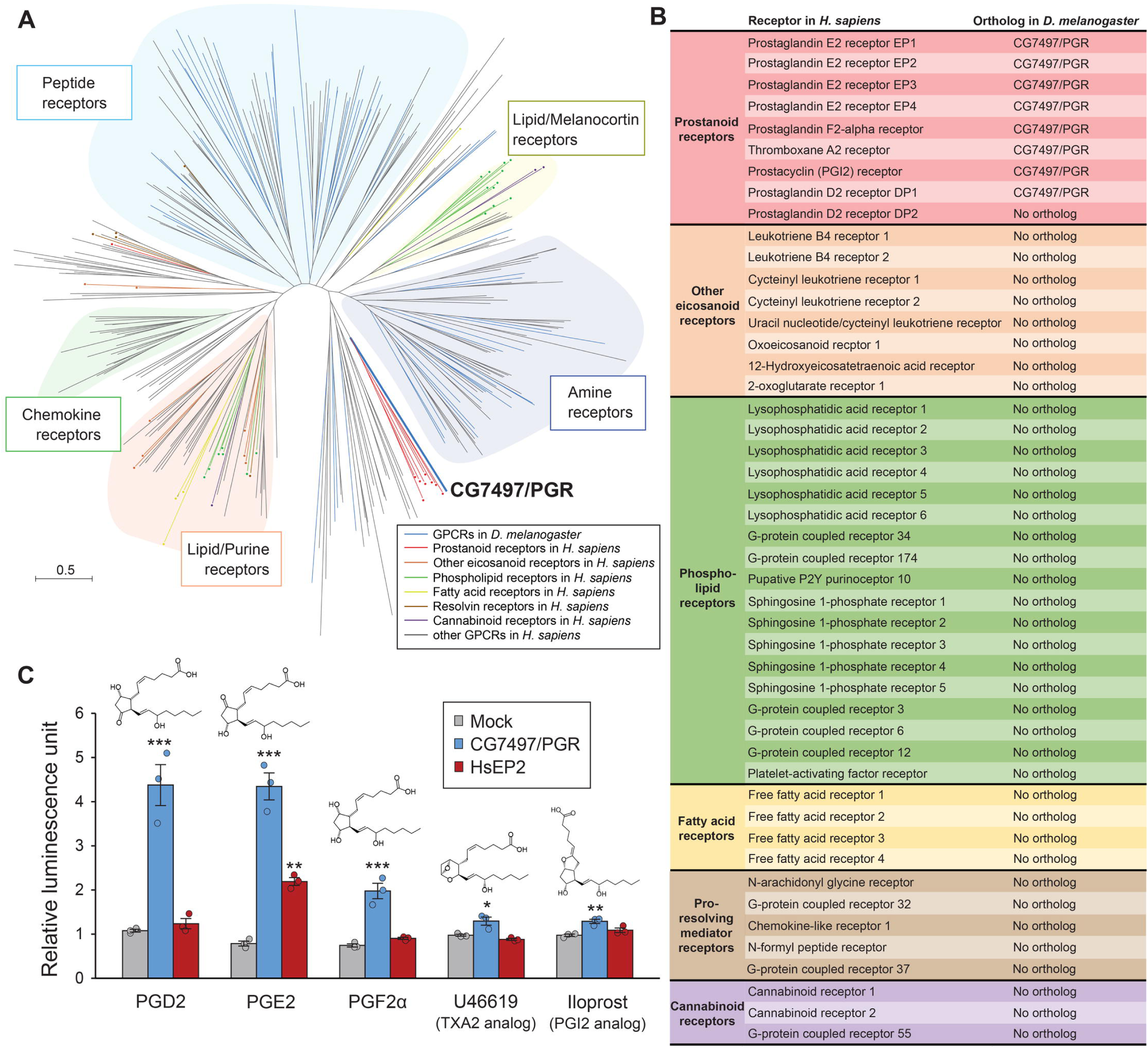
Identification of a prostanoid receptor in *D. melanogaster*. (A) Unrooted maximum-likelihood phylogenetic tree of class A G protein-coupled receptors (GPCRs) in *H. sapiens* and *D. melanogaster*. GPCRs in *D. melanogaster* are indicated by blue branches, whereas branches for *H. sapiens* GPCRs are color-coded based on different classes of ligands. The scale bar indicates an evolutionary distance of 0.5 amino acid substitutions per site. Opsins and leucine-rich repeat containing GPCRs are not included in the phylogenetic tree. Accession numbers of the receptors analyzed are listed in S1 Table. (B) List of lipid mediator receptors in *H. sapiens* and their *Drosophila* orthologs. (C) Luminescence responses of S2 cells co-expressing *Aequorin* and *Gα15*, along with *CG7497/PGR* or *H. sapiens* PGE2 receptor *EP2* (*HsEP2*), to prostanoids or their synthetic analogs. Relative luminescence normalized by luminescence in ligand-free controls is shown. Cells expressing *CG7497/PGR* were significantly activated by all ligands, whereas cells expressing *EP2* were activated only by PGE2. n = 3. \**p* < 0.05, \*\**p* < 0.01, \*\*\**p* < 0.001 (Dunnett’s test *vs* mock treatment control).

### PGR is required in tracheal cells for normal pupa-adult development

*PGR* expression pattern was visualized using *PGR-Gal4* generated by the CRISPR-mediated integration cassette system [29], where *T2A-Gal4* and 3xP3-GFP-encoding sequence were inserted in the *PGR* locus, so that Gal4 is expressed under the control of the endogenous *PGR* promoter (S2A Fig). *PGR* expression was detected in multiple tissues during *Drosophila* development (Figs 2A and S3). In adults, it is expressed in the gonads, gut, dorsal vessel, and hemocytes, as well as in the tracheae (Fig 2B). In particular, *PGR* is expressed in the tracheae in all developmental stages tested, suggesting its important functions in regulating development of the tracheal system. Indeed, *PGR* knockdown in the tracheae or terminal tracheal cells (TTCs) caused pharate adult lethality or eclosion defects (Fig 2B), suggesting that *PGR* expression in tracheal cells is necessary for normal progression of pupa-adult development.

**Fig 2.**
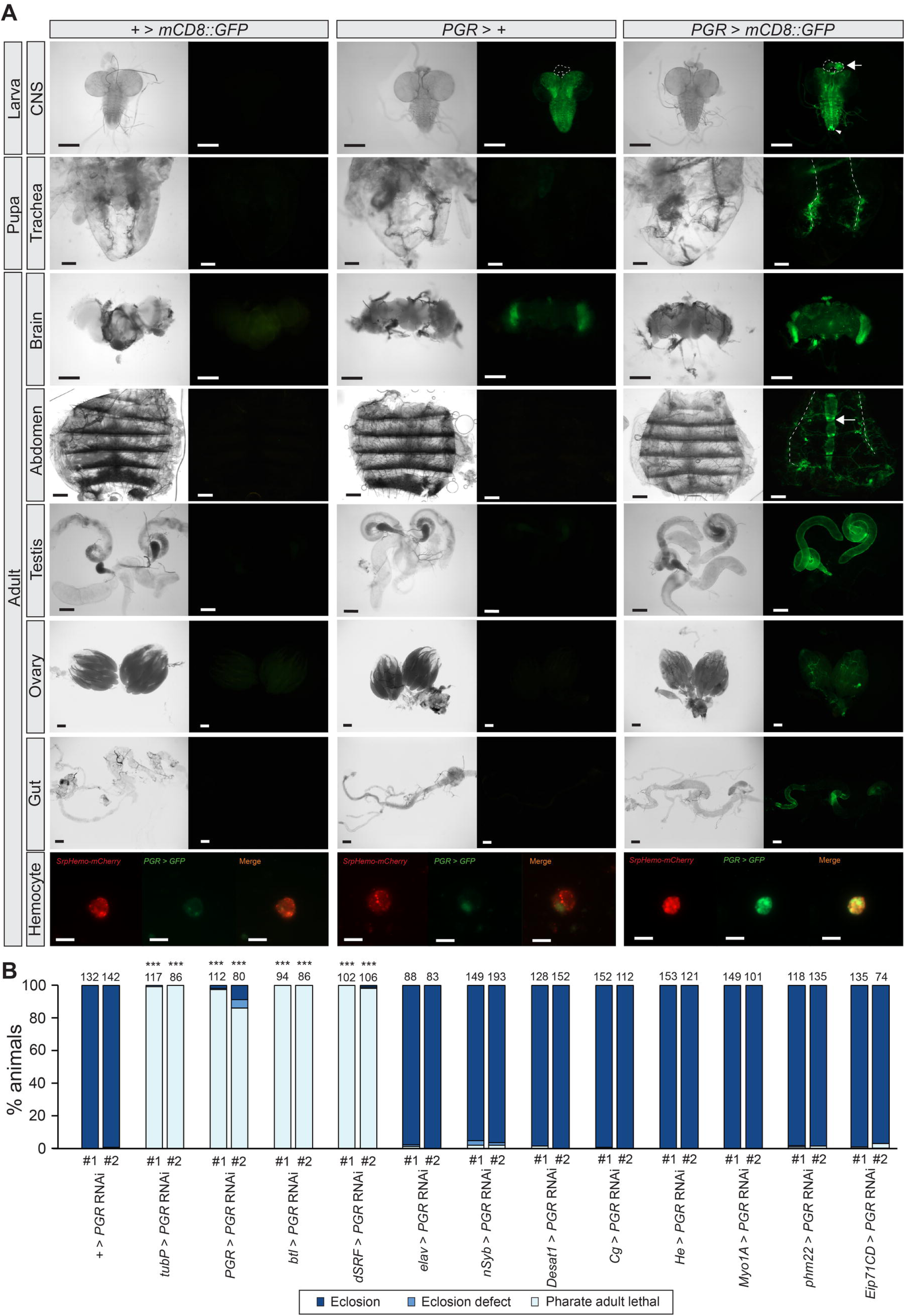
Expression of *PGR* during development. (A) Expression patterns of *CG7497*/*PGR* as visualized by *PGR*-*Gal4*-driven expression of *UAS-mCD8::GFP*. *PGR* is expressed in the prothoracic gland (outlined by the dashed line and marked by an arrow) and neurons located on the posterior end of the ventral nerve cord (indicated by an arrowhead) in third instar larvae. *PGR* is also expressed in the pupal and adult tracheae (dorsal trunks are indicated by dashed lines), as well as in testes, ovaries, dorsal vessel (indicated by an arrow), gut, and hemocytes in adults. Pupae are 48 hours after puparium formation, and adults are 72 hours after eclosion. Note the strong background GFP signals in the larval central nervous system (CNS) and adult optic lobes due to the *3xP3-GFP* marker in the *PGR*-*Gal4* line. Systemic GFP expression at different developmental stages is shown in S3 Fig. (B) Developmental phenotype caused by two independent *PGR* RNAi lines. *PGR* knockdown caused lethal phenotype at the pharate adult stage when induced either ubiquitously *(tubP-Gal4*), in *PGR* expression sites (*PGR-Gal4*), in tracheal cells (*btl-Gal4*), or in terminal tracheal cells (*dSRF-Gal4*). Numbers above the bars indicate flies analyzed for each genotype. Expression patterns of *Gal4* drivers are described in S11 Table. \*\*\**p* < 0.001 (Chi-square test *vs + > PGR* RNAi with Bonferroni correction). Scale bars, 200 µm; 10 µm in the hemocyte images.

### PGR promotes adult tracheogenesis

We next generated *PGR* knockout flies using the CRISPR-Cas9 system. Two mutant strains generated by independent guide RNA (gRNA) pairs are expected to be null mutants, as their exons are almost completely deleted (Fig 3A). Homozygous and transheterozygous *PGR* mutants died during late pupa-adult development after wing pigmentation, similar to its trachea-specific knockdown flies (Figs 3B and 3C). The lethal phenotype was significantly rescued by restoring *PGR* expression in TTCs using *dSRF*-*Gal4* (Fig 3D). Although *PGR* overexpression in all tracheal cells by *btl-Gal4* did not rescue the lethality, more adult flies eclosed when *PGR* was overexpressed using both *btl-Gal4* and *dSRF-Gal4* in the transheterozygous mutant background.

**Fig 3.**
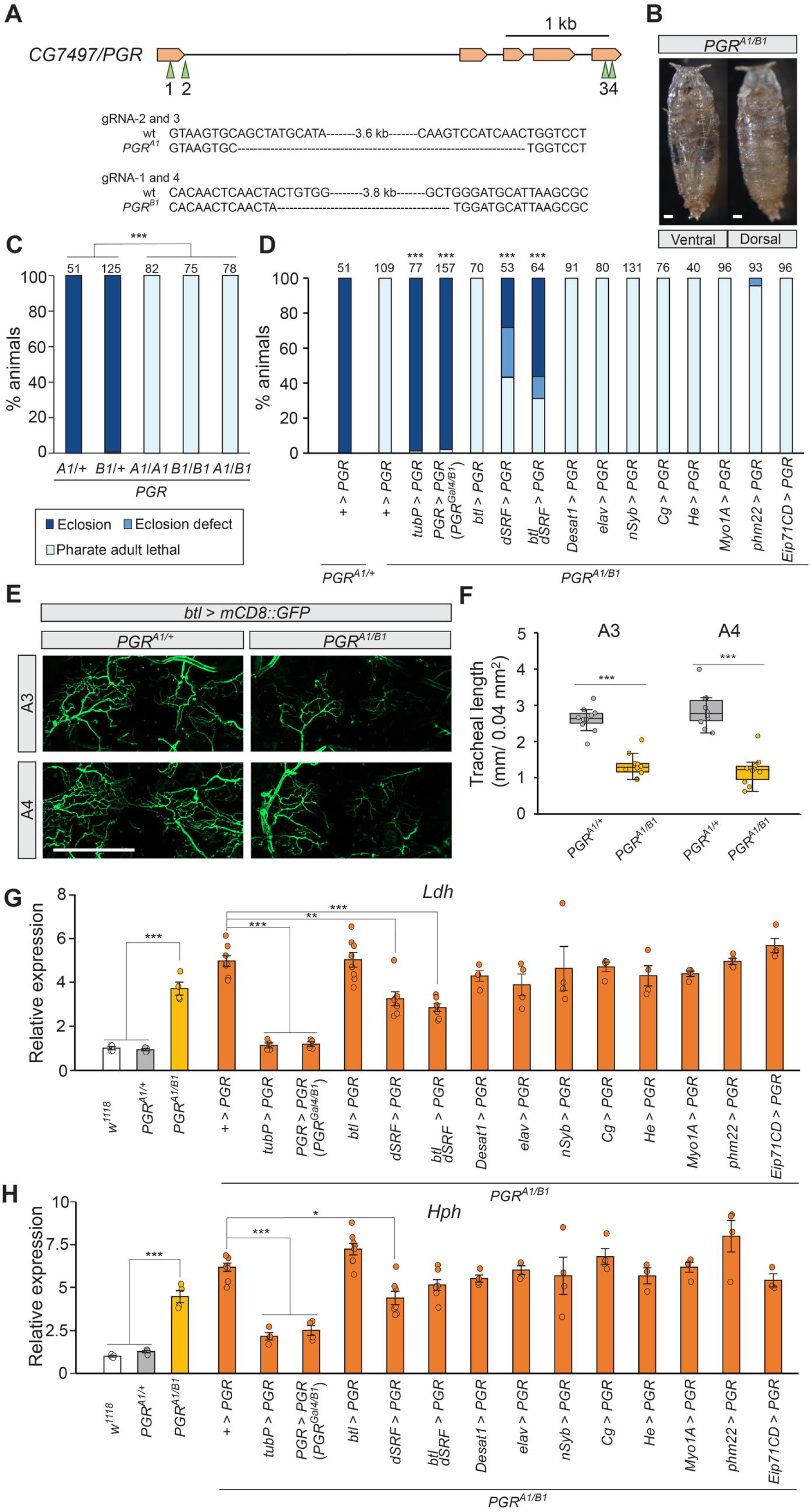
Phenotypic analysis of *CG7497/PGR* mutants. (A) Schematic diagram of target sites of mutation on the *CG7497*/*PGR* locus. Two mutant alleles were generated using two pairs of different gRNAs. Genomic sequences of the mutants are shown in S1 Document. (B) Representative images of a developmentally arrested transheterozygous *PGR* mutant at the pharate adult stage. (C) Developmental phenotype of the *PGR* mutants. Most homozygous and transheterozygous *PGR* knockout flies died as pharate adults. \*\*\**p* < 0.001 (multiple comparison Chi-square test with Bonferroni correction). Numbers above the bars indicate flies analyzed in each genotype. (D) Rescue of transheterozygous *PGR* mutants by *Gal4*-driven expression of *UAS-PGR*. The pharate adult lethality was rescued by overexpressing *PGR* either ubiquitously (*tubP-Gal4*), in *PGR* expression sites (*PGR-Gal4*), or in TTCs (*dSRF-Gal4*). \*\*\**p* < 0.001 (Chi-square test *vs + > PGR* with Bonferroni correction). As *PGR-Gal4* is a loss of function allele of *PGR*, the transheterozygous mutant of *PGR^Gal4^*/*PGR^B1^* was used when restoring *PGR* expression using *PGR-Gal4*. Numbers above the bars indicate flies analyzed for each genotype. (E) The tracheae were visualized by *btl-Gal4*-driven expression of *UAS-mCD8::GFP* in the third (A3) and fourth (A4) abdominal segments at 72 hours after puparium formation (APF). (F) Total tracheal length in the third (A3) and fourth (A4) abdominal segments in selected areas. The transheterozygous *PGR* mutant showed defective tracheal development in both segments. n = 10. \*\*\**p* < 0.001 (Student’s *t*-test). (G, H) Relative expression levels of hypoxia response genes, *Lactate dehydrogenase* (*Ldh*) (G) and *HIF prolyl hydroxylase* (*Hph*) (H), in *PGR* mutants at 72 hours APF. Transheterozygous *PGR* mutants showed higher expression of the hypoxia response genes. High expression of the hypoxia response genes in *PGR* mutants was suppressed by expressing *PGR* either ubiquitously (*tubP-Gal4*), in *PGR* expression sites (*PGR-Gal4*), or in TTCs (*dSRF-Gal4*). Expression levels are normalized by the levels of a reference gene, *rp49*, in the same cDNA samples and shown as relative to *PGR^A1/+^*. n = 3–8. \*\*\**p* < 0.001 (among *w^1118^*, *PGR^A1/+^* and *PGR^A1/B1^*, Tukey’s honestly significant difference test). \**p* < 0.05, \*\**p* < 0.01, \*\*\**p* < 0.001 (Rescue by *PGR* overexpression, Dunnett’s test *vs PGR^A1/+^*). Scale bars, 200 µm in B and E.

These results suggest that *PGR* expression in tracheal cells, particularly in TTCs, is necessary for normal development. Consistent with this, *PGR* knockout caused underdevelopment of the adult tracheal system in the abdomen (Figs 3E and 3F). Length of tracheoles in the *PGR* mutants was significantly shorter than control, indicating that PGR is necessary for promoting adult tracheal development. Similar defects in tracheal development were observed when the tracheae were visualized by GFP using another tracheal driver, *Trh-Gal4* (S4A and S4B Figs). Moreover, overexpression of *PGR* in the tracheae and TTCs using *btl-Gal4* and *dSRF-Gal4* significantly rescued underdeveloped tracheoles in *PGR* mutants (S4C and S4D Figs). Expression of hypoxia response genes, *Lactate dehydrogenase* (*Ldh*) and *HIF prolyl hydroxylase* (*Hph*), was upregulated in the *PGR* mutants, consistent with their defective tracheal development (Figs 3G and 3H). The high expression of the hypoxia response genes in the *PGR* mutants was suppressed by restoring *PGR* expression in TTCs (Figs 3G and 3H), in line with the developmental rescue experiments. Altogether, these results indicate that PGR regulates adult tracheal development, which is necessary for sufficient oxygen supply during pupa-adult development.

### PGR promotes tracheogenesis during early pupal stages to provide oxygen required for pupa-adult development

In order to determine the exact timing of PGR requirement in adult tracheal development, detailed expression analysis of hypoxia response genes in the *PGR* mutants was conducted (Fig 4A and 4B). Expression of both *Ldh* and *Hph* in the *PGR* mutants was higher than control flies beginning at 16 hours after puparium formation (APF). Since pupation happens approximately at 12 hours APF, PGR seems to function immediately after pupation to promote adult tracheal development. To further confirm this, a temporal knockdown experiment using a temperature-sensitive Gal4 suppressor, Gal80^ts^, was conducted (Fig 4C). At 18°C, flies pupate at about 24 hours APF. When *PGR* knockdown in the tracheae was initiated by 30 hours APF by transferring animals from 18°C to 29°C, most flies died before eclosion. However, the lethality gradually decreased as the onset of knockdown was shifted up to 48 hours APF, after which no significant pupal lethality was observed. Likewise, in a temporal rescue experiment of the *PGR* mutants by overexpressing *PGR*, most flies eclosed when the overexpression was initiated before 24 hours APF. However, the lethality was increased as the onset of overexpression was shifted up to 57 hours APF, after which no significant rescue happened. These results indicate that PGR is required for tracheogenesis during early stages of pupa-adult development after pupation.

**Fig 4.**
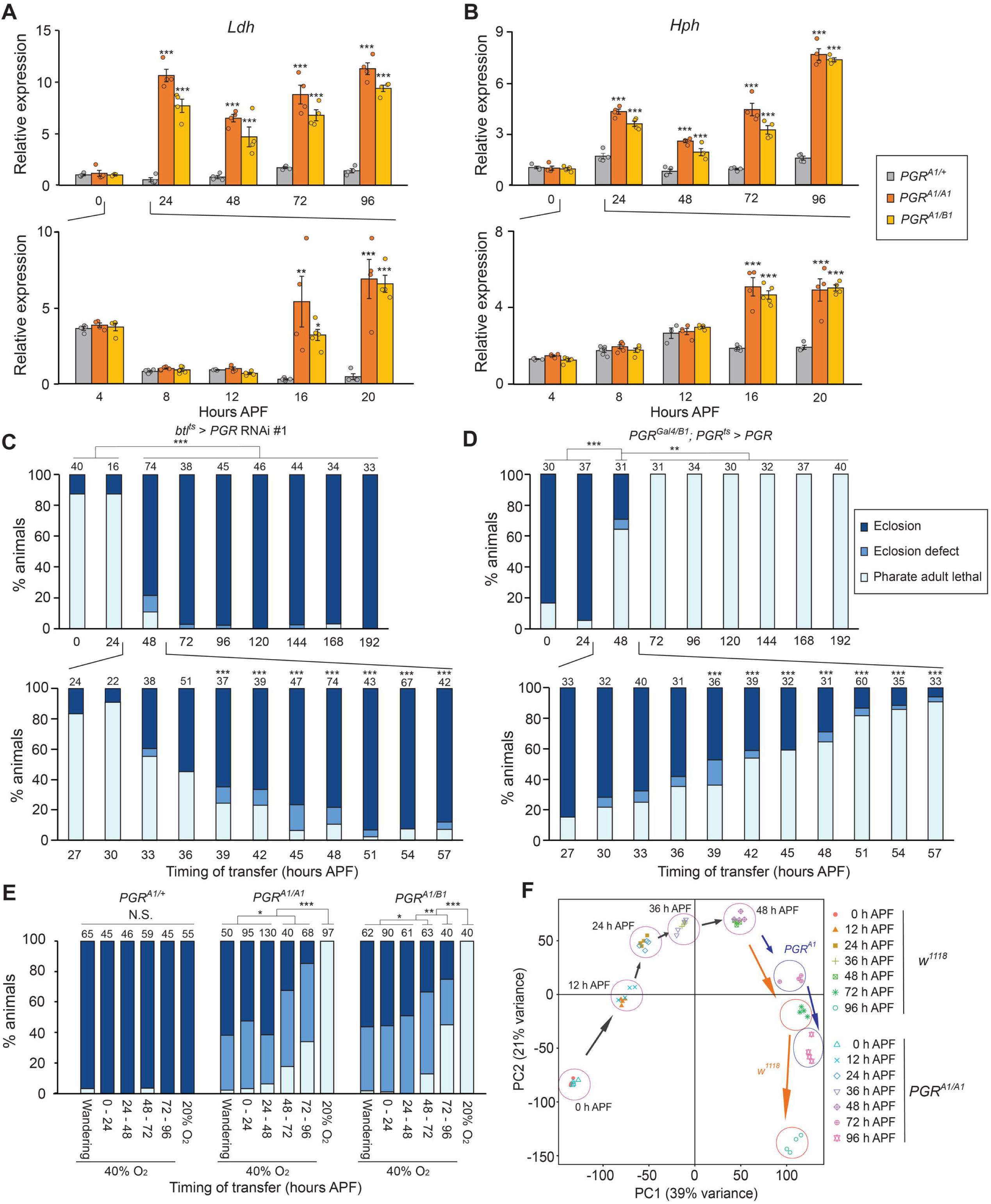
Stage-specific requirement of *PGR* during pupa-adult development. (A, B) Relative expression levels of hypoxia response genes, *Ldh* (A) and *Hph* (B), in *PGR* mutants between 0 and 96 hours after puparium formation (APF). Insects pupated about 12 hours APF. Homozygous and transheterozygous *PGR* mutants showed higher expression of both genes as compared to the heterozygous control after 16 hours APF. Expression levels are normalized by the levels of a reference gene, *rp49*, in the same cDNA samples and shown as relative to *PGR^A1/+^* at 0 hours APF. n = 3–6. \*\**p* < 0.01, \*\*\**p* < 0.001 (Dunnett’s test *vs PGR^A1/+^*). (C) Developmental deficiency caused by *PGR* RNAi using *btl-Gal4* and *tubP-Gal80^ts^*. Insects were reared at 18°C and transferred to 29°C at indicated hours APF. Flies pupated at ∼24 hours APF at 18°C. \**p* < 0.05, \*\*\**p* < 0.001 (upper panel, multiple comparison Chi-square test with Bonferroni correction; lower panel, Chi-square test *vs* 24 hours APF with Bonferroni correction). Numbers above the bars indicate flies analyzed for each treatment. (D) Rescue of the transheterozygous *PGR* mutant by *PGR*-*Gal4*-driven temporal expression of *UAS-PGR*. Insects were reared at 18°C and transferred to 29°C at indicated hours APF. \*\**p* < 0.01, \*\*\**p* < 0.001 (upper panel, multiple comparison Chi-square test with Bonferroni correction; lower panel, Chi-square test *vs* 24 hours APF with Bonferroni correction). Numbers above the bars indicate flies analyzed for each treatment. (E) Rescue of *PGR* mutants by high oxygen condition. Animals were kept in a container with 40% O_2_ from indicated stages until eclosion. \**p* < 0.05, \*\**p* < 0.01, \*\*\**p* < 0.001 (multiple comparison Chi-square test with Bonferroni correction). Numbers above the bars indicate flies analyzed for each treatment. (F) Principal component analysis based on whole body transcriptome of *w^1118^*and *PGR^A1/A1^*at selected hours APF. mRNA components showed similar patterns between *w^1118^* and *PGR^A1/A1^* from 0 hours until 48 hours APF, whereas different patterns were observed thereafter.

As the *PGR* mutants exhibited the hypoxia response due to defects in adult tracheal development, we reasoned that their developmental defects can be recovered by high concentrations of oxygen. We therefore transferred the *PGR* mutants to a growth chamber containing 40% oxygen at different stages of development (Fig 4E). More than half of the mutants eclosed when they were transferred to 40% oxygen by the mid pupal stage. Some flies eclosed even when they were transferred to 40% oxygen during late pupa-adult development, although the rescue rates were lower. This result confirmed that the *PGR* mutants die due to insufficient oxygen supply caused by defective tracheogenesis during pupa-adult development. Under normal oxygen concentration, the *PGR* mutants show no discernible developmental defects up to the mid pupal stage and eventually die at the pharate adult stage. Consistent with this observation, the whole-body transcriptome analysis revealed that gene expression patterns in control and the *PGR* mutants are similar until mid pupa-adult development (48 h APF; Fig 4F). In contrast, the *PGR* mutants showed distinct gene expression patterns at 72 and 96 hours APF. Overall, these results suggest that PGR functions between pupation and mid pupa-adult development to induce adult tracheal development, whose deficiency causes developmental defects in late pupal stages due to insufficient oxygen supply and leads to pharate adult lethality.

### Prostaglandins are synthesized in *D. melanogaster*

What are the endogenous ligands for PGR that promote adult tracheogenesis in *D. melanogaster*? In mammals, multiple active eicosanoids are synthesized from AA (Fig 5A). Although there is no obvious cyclooxygenase (PGG/H synthase or COX) ortholog among ten heme peroxidases conserved in insects [30] (S5 Fig, S3 Table), one heme peroxidase has been proposed to function as COX in *D. melanogaster* [11]. Moreover, based on our phylogenetic analyses, PGD, PGE, and PGF synthases are highly conserved in insects, although *D. melanogaster* lacks some of them (Figs 5B and S6–S10, S4–S8 Tables). On the other hand, among multiple cytochrome P450 enzymes in insects, there are no obvious orthologous enzymes of CYP5A1 (thromboxane A synthase) or CYP8A1 (prostacyclin synthase) (S11 Fig, S9 Table). These results suggest that prostaglandins are the major eicosanoids in insects. Indeed, PGD, PGE, and PGF have been detected in some insects [31,32], including *D. melanogaster* [22]. Since PGR signaling in tracheogenesis is critically required during early pupal stages, we reasoned that active PGR ligands are synthesized in the same time window during *Drosophila* development. Therefore, Eicosanoid contents were analyzed in the whole-body extract of early pupae by liquid chromatography-tandem mass spectrometry (LC-MS/MS). Under normal rearing conditions, ∼0.13 ng of AA was detected per animal, whereas prostanoids were undetectable. In contrast, when larvae were reared on the food supplemented with AA, significant amounts of PGD2, PGE2, and PGF2α were detected, whereas no other prostanoids were observed (Fig 5C). These results indicate that flies have the ability to synthesize PGD, PGE, and PGF, consistent with the presence of multiple prostaglandin synthase orthologs. Considering the high activity of PGD2 and PGE2 as PGR ligands (Fig 1C), it is conceivable that either PGD2 or PGE2 is an endogenous PGR ligand that promotes tracheogenesis during pupa-adult development.

**Fig 5.**
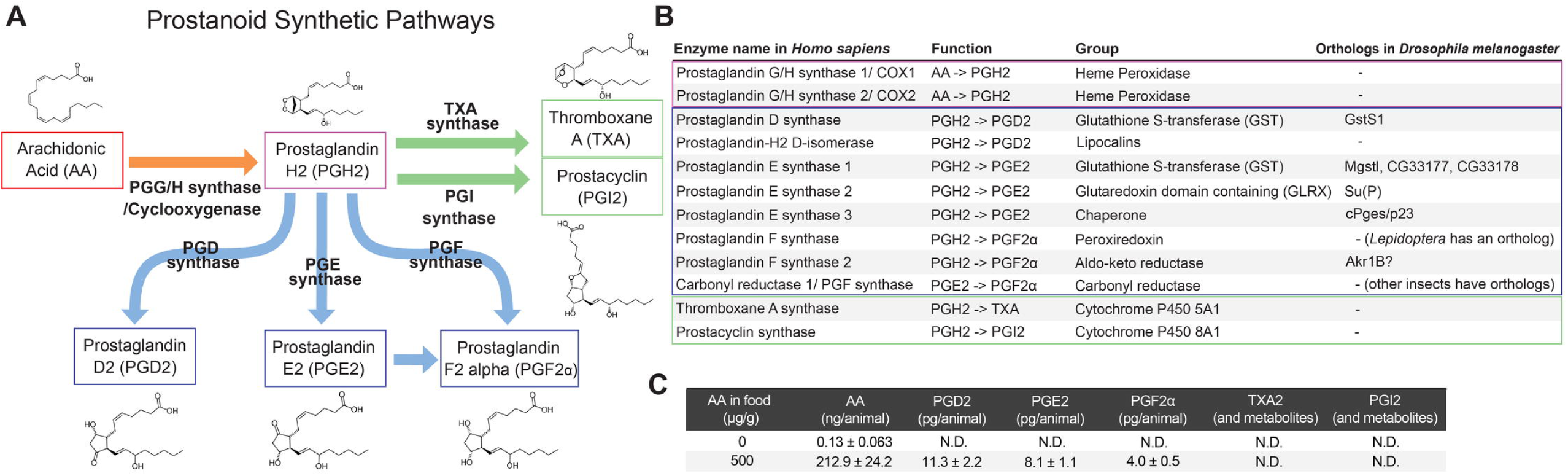
Conserved prostanoid synthases in *D. melanogaster*. (A) Schematic diagram of prostanoid synthetic pathways. Multiple prostanoids are synthesized from arachidonic acid (AA) by different enzymes. (B) *Drosophila* enzymes orthologous to human prostanoid synthases were identified by phylogenetic analyses. Phylogenetic trees are shown in S5–S11 Figs. PGD, PGE, and PGF synthases are highly conserved in insects, although *D. melanogaster* lacks some of them. (C) Prostanoids detected in whole body extracts of pupae by liquid chromatography-tandem mass spectrometry (LC-MS/MS)-based lipidomics. AA was detected in intact pupae at 0–4 hours after pupation, whereas prostanoids were undetectable. PGD2, PGE2, and PGF2α were detected in pupae that were raised in AA-supplemented diet during larval development. Data shown are mean total amount of eicosanoids in the whole body per animal ± SEM. n = 4–5.

### Cytosolic Prostaglandin E synthase/p23 activates local prostaglandin signaling to promote tracheogenesis

In order to identify prostaglandin synthase(s) that are responsible for tracheal development, we generated null mutants of six *Drosophila* orthologs of PGD and PGE synthases (S12 Fig). Among these mutants, a triple knockout mutant of three paralogous genes, *Mgstl*, *CG33177*, and *CG33178* (*Prostaglandin E synthase 1* or *PTGES1* orthologs) and a mutant of *cytosolic Prostaglandin E synthase (cPges)/p23* (a *PTGES3* ortholog) showed pupal lethal phenotype (Fig 6A). In particular, most *cPges/p23* mutant animals died during pupa-adult development, although more than half of them died without pigmentation (Fig 6B). Importantly, only *cPges/p23* mutant pupae showed the hypoxia response after 24 hours APF (Figs 6C and 6D) similar to the *PGR* mutants (Figs 4A and 4B), suggesting that cPges/p23 is the major PGE synthase that produces PGR ligands during early pupa-adult development. It should be noted that cPges/p23, as well as PTGES3 in vertebrates, functions as a co-chaperone. In mammals, PTGES3 acts as a component of the heat shock protein 90 (Hsp90)-based chaperon complex and facilitates functions of its target proteins such as the estrogen receptor [33]. *Drosophila* cPges/p23 also stabilizes Hsp90 by acting as a co-chaperon [34]. This potentially explains the high lethality of the *cPges/p23* mutant pupae, as Hsp90 has important functions in *Drosophila* development [35].

**Fig 6.**
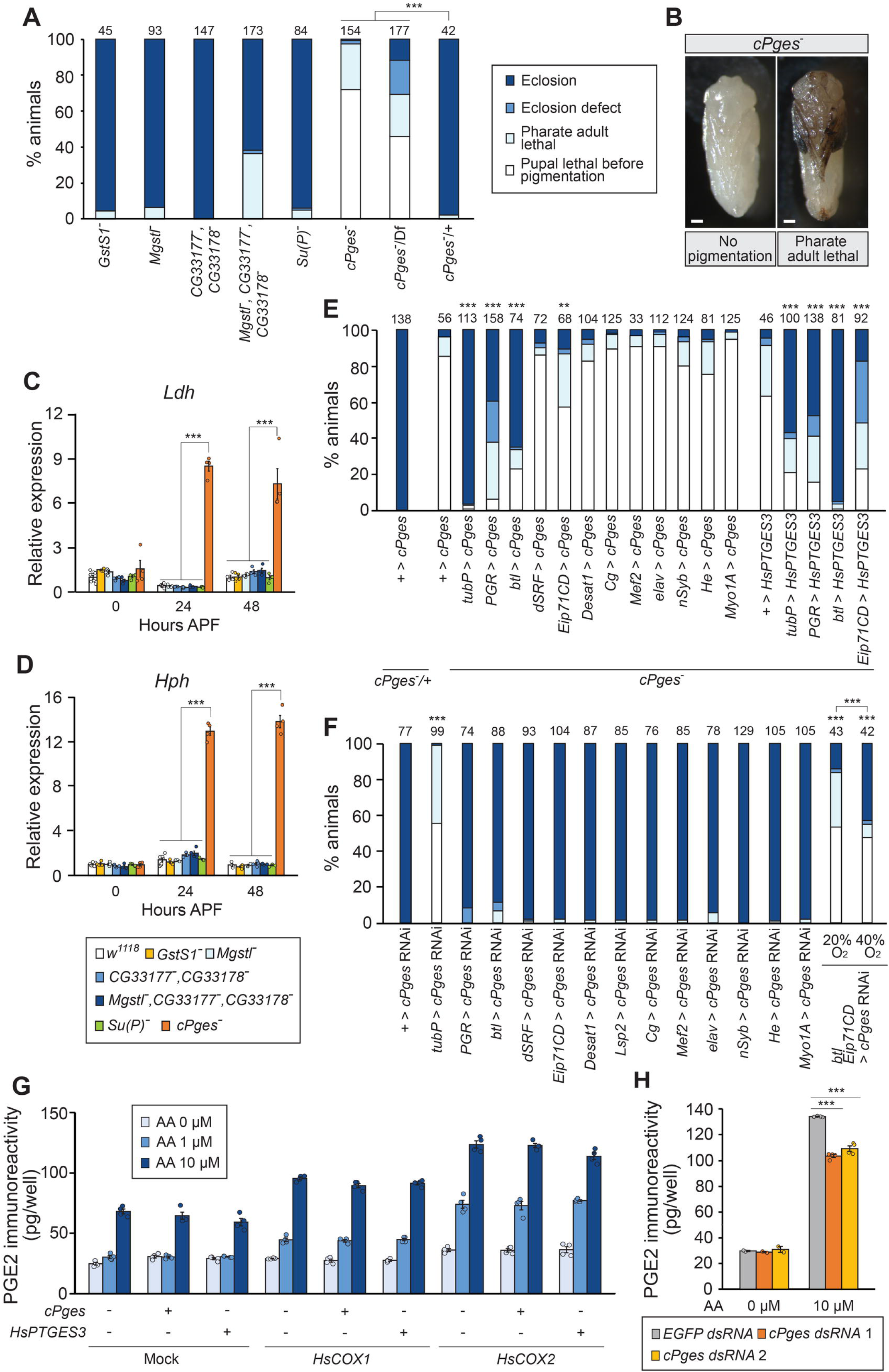
Functional characterization of a *Drosophila* PGE synthase. (A) Developmental phenotype of PGD and PGE synthase ortholog mutants. Null mutant of *cPges*/*p23* and triple mutants of *Mgst^-^*, *CG33177*, and *CG33178* died during pupa-adult development. Multiple comparison Chi-square test with Bonferroni correction was applied among *cPges*/*p23* mutants including the heterozygous control. \*\*\**p* < 0.001. (B) Representative images of *cPges*/*p23* knockout flies. About 70% of flies died before wing pigmentation, whereas ∼30% of flies died as pharate adults. Scale bar, 200 µm. (C, D) Relative expression levels of hypoxia response genes, *Ldh* (C) and *Hph* (D), in the PGD and PGE synthase ortholog mutants. The *cPges*/*p23* mutant showed high expression of hypoxia response genes at 24 and 48 hours after puparium formation (APF). Expression levels are normalized by the levels of a reference gene, *rp49*, in the same cDNA samples and shown as relative to 0 hours APF of *GstS1^-^*. n = 3–8. *p* < 0.001 (Tukey’s honestly significant difference test). (E) Rescue of the *cPges*/*p23* mutant by *Gal4*-driven expression of either *cPges*/*p23* or human *prostaglandin E synthase 3* (*HsPTGES3*). The lethal phenotype was rescued by overexpressing *cPges*/*p23* or *HsPTGES3* either ubiquitously (*tubP-Gal4*), in *PGR* expression sites (*PGR-Gal4*), in tracheal cells (*btl-Gal4*), or in the epidermis (*Eip71CD-Gal4*). \*\**p* < 0.01, \*\*\**p* < 0.001 (Chi-square test *vs + > cPges*/*p23* or *+ > HsPTGES3* with Bonferroni correction). (F) Developmental deficiency caused by *cPges*/*p23* RNAi. *cPges*/*p23* knockdown flies died during pupa-adult development when induced using either a ubiquitous driver *(tubP-Gal4*) or a combination of the tracheal cell (*btl-Gal4*) and epidermis (*Eip71CD-Gal4*) drivers. Lethality caused by *cPges*/*p23* RNAi in tracheal cells and epidermis was significantly rescued by 40% O_2_. \*\*\**p* < 0.001 (multiple comparison Chi-square test *vs + > cPges*/*p23* RNAi with Bonferroni correction, or 20% O_2_ *vs* 40% O_2_). (G) PGE2 conversion assay using *Drosophila* S2 cells. Arachidonic acid (AA) was added to the culture medium of S2 cells transfected with *cPges*/*p23* or *HsPTGES3* along with a *H. sapiens cyclooxygenase* (*HsCOX1* or *HsCOX2*), and PGE2 amount in the medium was determined. The basal levels of PGE2 immunoreactivity in all samples is likely due to the cross-reactivity of the antibody used in the assay. Increased PGE2 immunoreactivity was detected in *cyclooxygenase*-expressing cells regardless of *cPges*/*p23* or *HsPTGES3* transfection. n = 4. (H) Reduction of the PGE2 synthetic activity of S2 cells by knocking down endogenous *cPges*/*p23* expression. AA was added to the culture medium of S2 cells transfected with *HsCOX2* and treated with *cPges*/*p23* dsRNAs. PGE2 amount in the medium was determined. n = 2 (AA 0 µM) or n = 4 (AA 10 µM). \*\*\**p* < 0.001 (Dunnett’s test *vs EGFP* dsRNA-treated cells).

Tissue-specific overexpression of *cPges/p23* using either *PGR-Gal4*, *btl-Gal4*, or an epidermis driver *Eip71CD-Gal4* rescued the lethal phenotype of the *cPges/p23* mutant (Fig 6E). Consistent with this, *cPges/p23* knockdown using a combination of *btl-Gal4* and *Eip71CD-Gal4* drivers caused high lethality, which was significantly rescued by high concentrations of oxygen (Fig 6F). These results suggest that prostaglandins are synthesized in the epidermis and trachea to activate PGR in TTCs, thus forming a local prostaglandin signaling network that promotes tracheogenesis during early pupal stages.

In order to confirm that cPges/p23 functions as a PGE synthase, we performed an enzymatic conversion assay using *Drosophila* S2 cells. As an intermediate of PGE2 synthesis, prostaglandin H2 (PGH2), is unstable, human COXs were co-expressed with either *cPges/p23* or human *PTGES3* to detect PGE2 synthesis from AA using the enzyme-linked immunosorbent assay (ELISA) that is highly specific to PGE2 (S13A Fig). Interestingly, PGE2 synthesis was detected without *cPges/p23* or human *PTGES3* transfection, suggesting high endogenous PGE2 synthase activities in S2 cells (Fig 6G). Therefore, we applied double-stranded RNA (dsRNA) against *cPges/p23* to reduce expression of endogenous *cPges/p23* (S13B Fig). As a result, PGE2 production was significantly suppressed by *cPges/p23* knockdown, indicating that cPges/p23 functions as a PGE synthase in *D. melanogaster* (Fig 6H).

## Discussion

In this study, we revealed that a single GPCR that we named PGR mediates prostaglandin signaling essential for adult tracheal development in *D. melanogaster*. PGE2 is synthesized in the trachea and epidermis by cPGES3/p23 and activates PGR in TTCs after pupation. This PGE2-PGR signaling induces tracheal development, which provides sufficient oxygen supply during pupa-adult development.

In mammals, there are numerous species of lipid mediators [3], which act through various GPCRs as shown in Fig 1. In contrast, our phylogenetic analyses clearly indicate that, among these lipid mediator GPCRs, only prostanoid receptors are conserved between mammals and insects, suggesting evolutionarily conserved functions of prostanoids in the animal kingdom. Among nine prostanoid receptors in *H. sapiens*, there is a considerable overlap of their cognate ligands, as well as a significant amount of crosstalk between their downstream signaling pathways [2]. Humans and other mammals thus have a highly complex prostanoid signaling network. In contrast, although PGD2, PGE2, and PGF2α have been detected in multiple insect species [22,31,32], our current study indicates that all of them likely act through a single receptor, PGR. Alternatively, it is also possible that there are as-yet-unknown insect prostanoid receptors that are not homologous to human lipid mediator receptors. Nonetheless, our comprehensive phylogenetic analyses of prostanoid biosynthetic enzymes indicate that prostaglandins are the primary, and potentially the only, prostanoids produced in insects (Fig 5). Altogether, prostaglandin-PGR signaling represents the primary lipid mediator signaling that is highly conserved in the animal kingdom, and our current study provides a solid basis as well as useful genetic tools for using *D. melanogaster* as an extremely simple model system to investigate this highly conserved signaling pathway.

Our phylogenetic study suggested that one PGD synthase and five PGE synthases are conserved in *D. melanogaster*. To date, several prostaglandin synthetic enzymes have been reported in insects [11,16,17]. A recent study in *D. melanogaster* suggested that cPges/p23 functions as a PGE synthase in the ovary [19], and our current study further confirmed its PGE synthetic function. During pupa-adult development, cPGES/p23 synthesizes PGE2 in the epidermis and tracheae to activate local PGR signaling, which is consistent with observations in mammals that prostanoids act locally at the site of their production [2]. It is likely that this local, paracrine nature of prostaglandin signaling keeps the whole-body prostaglandin titer below the detection limit, unless extra AA is provided in the food (Fig 5C). When larvae were fed with the AA-supplemented diet, we also detected PGD2 and PGF2α, as well as PGE2, in early pupae. Our study thus confirms that *D. melanogaster* has the ability to synthesize PGD and PGF, which is consistent with the presence of a PGD synthase ortholog (GstS1) in *D. melanogaster* (Fig 5B).

Although we could not find obvious PGF synthase orthologs, a recent study suggests that Aldo-keto reductase 1B (Akr1B) functions as a *Drosophila* PGF synthase [19]. Functional characterization of these putative prostaglandin synthases, including PGE synthase orthologs other than cPGES/p23, is clearly warranted in future studies.

Although prostanoids are highly pleiotropic signaling molecules [2], most prostanoid studies in insects have focused on their functions in immunity and reproduction [9–19,21,22]. To our knowledge, this is the first study to demonstrate that prostaglandin signaling promotes tracheal development in insects. A fibroblast growth factor named branchless (bnl) and its receptor breathless (btl) have been well investigated as important signaling molecules for branching and guidance of the tracheae in *D. melanogaster* [36]. Growth of TTCs is controlled by oxygen demand, as hypoxia stimulates *bnl* expression in target tissues to promote tracheal outgrowth in both larvae and adults [37–39]. In contrast, however, it is reported that *bnl* expression is not upregulated under hypoxic conditions immediately after pupation [40]. Indeed, expression levels of *bnl* in heterozygous and homozygous *PGR* mutants were almost the same until the pharate adult stage (S14A Fig), even though homozygous *PGR* mutants exhibited hypoxia responses throughout the pupal stage (Figs 4A and 4B). These results suggest that oxygen demand does not activate bnl-btl signaling during the pupal stage, and prostaglandin signaling acts as an alternative tracheogenic signaling pathway during pupa-adult development. Interestingly, *PGR* mutant flies rescued by high oxygen supply no longer require high oxygen for their survival after eclosion. This is likely due to the elevated bnl-btl signaling induced by hypoxia immediately after eclosion (S14B Fig), which is largely suppressed within 24 hours (S14C Fig). Indeed, the underdeveloped tracheal system in *PGR* mutants rescued by 40% oxygen quickly recovers to the level of the control heterozygous mutants within 24 hours after eclosion (S14D Fig). Overall, these results suggest that oxygen demand activates bnl-btl signaling in adult flies as previously reported [36,39,40], which induces compensatory tracheogenesis in *PGR* mutant flies after eclosion.

Considering that PGR is the only evolutionarily conserved prostanoid receptor in the *Drosophila* genome, it is possible that PGR mediates all major prostanoid-related biological processes in flies. Consistent with this, our transcriptome analysis revealed lower expression of antimicrobial peptides and reproduction-related genes in *PGR* mutant prepupae and pharate adults, respectively (S10 Table, S15 Fig), suggesting potential involvement of prostaglandin signaling in immunity and reproduction. Results of our current study, along with the genetic tools we developed, are expected to provide valuable resources for future studies to investigate such pleiotropic functions of prostaglandin signaling beyond development, which may give us insights into this highly conserved signaling pathway from evolutionary perspectives.

## Materials and methods

### Flies

Flies were raised at 25°C under 12 h-light and 12 h-dark photoperiod. The animals were fed on standard cornmeal diet containing 6 g *Drosophila* agar type II (Genesee Scientific, #66-103), 100 g D-(+)-glucose (SIGMA, #G8270-25KG), 50 g inactive dry yeast (Genesee Scientific, #62-106), 70 g yellow cornmeal (Genesee Scientific, #62-101), 6 ml propionic acid (SIGMA, #402907-500ML), and 10 ml Tegosept (Genesee Scientific, #20-258) in 1,025 ml of water. The control strain *w^1118^* and transgenic flies were obtained from the Bloomington *Drosophila* Stock Center (BDSC) and Vienna *Drosophila* Resource Center (VDRC) as shown in S11 Table.

Vectors to construct *UAS-PGR* and *UAS-cPges*/*p23* fly lines were obtained from the *Drosophila* Genomic Resource Center (DGRC, UFO02753 and UFO02035; S12 Table). *pUAST-HsPTGES3* was prepared from *pCMV6-HsPTGES3* from ORIGENE and cloned into the *pUAST* vector. All new transgenic flies were generated by BestGene Inc. Gene deletion mutant flies were generated using the CRISPR-Cas9 system: *PGR^A1^* and *PGR^B1^* were generated as described below, whereas the other mutant strains were generated by WellGenetics Inc. using CRISPR-based homologous recombination. Schematic diagrams and detailed sequences of the mutants are shown in S12 Fig and S2-6 Documents, respectively. All mutants were analyzed after backcrossing with the *w^1118^*control strain at least 4 times to minimize potential effects of off-target mutations.

### Cell lines

S2 cells used for aequorin luminescence assay and enzyme assays were obtained from DGRC (S2-DRSC, stock number 181) and maintained in 75 cm^2^ flask (VWR, #10062-860) in Shields and Sang M3 insect medium (Sigma-Aldrich, #S3652-500ML) containing 10% Insect Medium Supplement (Sigma-Aldrich, #I7267-100ML), 10% heat-inactivated fatal bovine serum (Gibco, #10082147), and 1% penicillin-streptomycin solution (Thermo Fisher Scientific, #15140122). Cells were incubated in a humidified incubator at 25°C.

### Phylogenetic tree analysis

Unrooted maximum-likelihood phylogenetic trees were generated using MEGAX [41]. Amino acid sequences of class A GPCRs (except for opsins and leucine-rich repeat-containing GPCRs) and eicosanoid synthases in *H. sapiens* and *D. melanogaster* were selected using HUGO Gene Nomenclature Committee at the European Bioinformatics Institute (https://www.genenames.org/) and Flybase [42], respectively. Entire amino acid sequences of the receptors and enzymes in the silkmoth (*Bombyx mori*), western honey bee (*Apis mellifera*), red flour beetle (*Tribolium casteneum*), pea aphid (*Acrythosiphon pisum*), and Nevada termite (*Zootermopsis nevadensis*) were obtained from National Center for Biotechnology Information database [43] using full length amino acid sequences of *H. sapiens* and *D. melanogaster* proteins as queries. The protein names and GenBank accession numbers are listed in S1–9 Tables.

### Aequorin-based calcium mobilization assay in S2 cells

Vectors for exogenous expression of *Aequorin*, *Gα15*, *CG7497/PGR*, and *HsEP2* in S2 cells were generated from *pLV-CMV-aequorin* (VectorBuilder), *pCMV6-GNA15* (Genomics-online), GH27361 (DGRC), and *pCMV6-HsEP2* (ORIGENE), respectively, and cloned into the *pBRAcPA* vector. S2 cells at a density of 2 million cells in 4 mL of the culture medium were seeded on a 60 mm petri dish (Falcon). Transfection of 1 µg of *pBRAcPA-Aequorin*, 2 µg of *pBRAcPA-Gα15*, and 1 µg of *pBRAcPA* containing either *CG7497/PGR* or *HsEP2* was performed 3 hours after seeding using Effectene Transfection Reagent (Qiagen, #301425) following the manufacturer’s protocol. After 3 days, 4.5 million cells in 2 mL of the assay medium (Shields and Sang M3 insect medium containing 10% Insect Medium Supplement) were transferred into a 6-well clear flat bottom multiple well plate (Corning). The cells were then incubated with 5 µM of coelenterazine-h (Promega, #S2001) for 4 hours with gently shaking.

After adding 1 ml of the assay medium, 50 µl of the cells (approximately 7,500 cells) were applied to a LUMIstar Omega microplate reader (BMG Labtech). In this system, the cells were injected into 50 µl of the assay medium containing 200 µM of a prostaglandin or a prostanoid analog (Cayman Chemical, PGD2, #12010; PGE2, #14010; PGF2α, #16010; U-46619, #16450; iloprost, #18215). Luminescence was detected every 0.2 seconds from 5 seconds before injection until 85 seconds after injection. Peak areas were calculated and normalized by areas obtained with no ligand containing medium.

### Visualization of *UAS-GFP*-expressing tissues

Developmental stages of flies were determined based on hours after egg laying (larvae), puparium formation (pupae), or eclosion (adults). Tissues were carefully dissected in PBS. *PGR* expressing tissues were visualized with *PGR-Gal4*-driven expression of *UAS-mCD8::GFP* [29]. GFP signals in the whole body and dissected tissues were observed using a SteREO Discovery.V12 microscope (Zeiss). GFP signal in hemocytes and the adult tracheal system were observed using a Zeiss Axio Imager M2 equipped with ApoTome.2 (Zeiss). SrpHemo-3xmCherry was used as a marker for hemocytes [44]. Pupae at 72 hours APF were dissected for observing dorsal tracheae [45]. Dissected dorsal tracheae were visualized by *btl-Gal4* or *Trh-Gal4*-driven expression of *UAS-mCD8::GFP*. For the PGR rescue experiment (S4C and S4D Figs), *UAS-mCD8::GFP* was driven by the combination of *SRF-Gal4* and *btl-Gal4*. Two 0.2 mm × 0.2 mm areas in the third and fourth abdominal segments were selected in each image. Tracheae were traced and the total tracheal length in each image was measured using Fiji with NeuronJ plugin [46].

### Scoring of lethal stages

Early pupae (up to 24 h APF) were collected and kept in *Drosophila* narrow vials (Genesee Scientific) containing wet tissue at 25°C under 12 h-light and 12 h-dark photoperiod. Phenotypes of flies were scored at 120 h after APF as either eclosion, eclosion defect, pharate adult lethal, or pupal lethal without wing pigmentation (lethal during early to middle pupa-adult development).

### Generation of *PGR* mutants

*PGR* mutant alleles (*PGR^A1^* and *PGR^B1^*) were generated using the CRISPR-Cas9 system as previously reported [47] with slight modifications. Pairs of gRNA target sequences (20 bp) were designed as shown in Fig 2A. Annealed oligonucleotides containing 20-bp target sequences (S13 Table) were inserted into the *pBFv-U6.2* or *pBFv-U6.2B* vector provided by the National Institute of Genetics [48]. The fragment containing the *U6* promotor and first gRNA in *pBFv-U6.2* was ligated into *pBFv-U6.2B* containing the second gRNA. Injection of plasmids and generation of G1 mutant strains were performed by BestGene Inc. Genotyping of *PGR^A1^* and *PGR^B1^* was conducted by PCR with extracted genomic DNA of flies with designed primers listed in S13 Table. Genomic sequences of the mutants are shown in S1 Document.

### Total RNA extraction and quantitative reverse transcription (qRT)-PCR

Four pupae were collected at each developmental stage in 1.5 mL tubes. Total RNA from animals was extracted using TRIzol reagent (Invitrogen, #15596018) according to the manufacture’s protocol. Extracted RNA was further purified using the RNeasy mini kit (Qiagen, #74104) and treated with RNase-Free DNase Set (Qiagen, #79254) following the recommended protocols. cDNA was synthesized from 500 ng RNA with PrimeScript RT Master Mix (Takara Bio) and diluted in 4x volume of TE buffer (10 mM Tris-HCl, 1 mM ethylenediamine-tetraacetic acid, pH 8.0). Amounts of mRNA were quantified by qRT-PCR on CFX connect real-time PCR detection system (Bio-Rad) with iQ SYBR Green Supermix (Bio-Rad, #1708882) using specific primers listed in S13 Table. For absolute quantification of mRNAs, serial dilutions of *pGEM-T* plasmids (Promega, #A3600) containing coding sequences of the target genes were used as standards. Transcript levels were normalized by *rp49* levels in the same samples.

### Temporal *PGR* knockdown and overexpression

Flies were raised at 18°C, and white pupae were transferred into *Drosophila* narrow vials (Genesee Scientific) containing wet tissue. Flies were then transferred to 29°C at different stages and reared until eclosion or for 4 days after transfer.

### Rescue of *PGR* mutants by high oxygen condition

Prepupae and pupae were collected every 24 hours and kept in normal condition until selected developmental timing. Animals were then transferred into plastic boxes, and oxygen was supplied up to 40% every 24 hours. Lethal stages were scored after 120 hours APF as described above.

### Transcriptome analysis

Eggs of *PGR^A1^* and *w^1118^* were laid on grape juice plates. Newly hatched larvae were transferred to standard cornmeal diet with less than 50 larvae/vials. Males and females were separated during the 3rd instar stage, and heterozygous mutants were removed using GFP signals in balancer-carrying larvae. Flies were transferred into vials containing wet tissue at puparium formation and kept until selected hours APF. Four biological replicates for each genotype and developmental timing (3 males and 3 females per sample) were collected and frozen in liquid nitrogen. Total RNA was extracted with the TRIzol reagent and RNeasy mini kit as described above. RNA qualification, library preparation, and RNA-seq were performed by Novogene Inc. Briefly, RNA quality was analyzed using an Agilent Bioanalyzer 2100. mRNA enriched using oligo(dT) beads was fragmented randomly, and cDNA was synthesized using mRNA template and random hexamer primers. After sequencing adaptor ligation, libraries were analyzed with NovaSeq 6000 (Illumina) to obtain 150-cycle paired-end sequencing, which produced more than 17.1 million reads per sample. Adaptor sequence-containing reads and low-quality reads were removed with the fastp software [49]. Reads were then aligned to the *Drosophila* genome (BDGP6.46, Ensembl) [50] with Hisat2 [51]. The number of aligned reads on each gene region was counted with samtools [52] and featureCount [53]. Differential expression analysis and principal component analysis were performed with DESeq2 packages using iDEP0.96 (http://bioinformatics.sdstate.edu/idep92/) [54]. Transcripts meeting a cutoff of 2-fold difference in mRNA abundance and false discovery rate of < 5% were considered as differentially expressed genes.

### Prostanoid detection

Eggs of *w^1118^* were laid on grape juice plates as described above. Newly hatched larvae were transferred to cornmeal diet containing selected amounts of AA with less than 50 larvae/vials. Fifty early pupae (0–4 hours after pupation) were collected in sample tubes and kept at −80°C until analysis. Extraction and detection of eicosanoids were performed at the UCSD Lipidomics Core. Briefly, 100 µl of 10% methanol with an internal standard mix (Cayman Chemical) was added to each sample, and pupae were homogenized by a bead homogenizer. Homogenates were then purified by polymeric reverse phase columns (Phenomenex, 8B-S100-UBJ) with 50 µl of elution buffer (63% de-ionized water, 37% acetonitrile, and 0.02% Acedic Acid). Eicosanoid determination was performed by reverse phase-ultra-performance liquid chromatography-tandem mass spectrometry with ACQUITY UPLC System (Waters) and SCIEX 6500 triple quadrupole linear ion trap mass spectrometer (AB Sciex) as described previously [55].

### Synthesis of *cPges*/*p23* double-stranded RNA

PCR was carried out using primers containing T7 promotor sequences (listed in S13 Table) using a *cPges*/*p23* cDNA clone LD23532 (DGRC) as a template. RNA was synthesized using the MEGAscript T7 Transcription Kit (Ambion, AM1334), followed by incubation with DNase for 30 min at 37°C. After incubation at 95°C for 5 min, the corresponding RNA products were mixed and gradually cooled down to room temperature for annealing. Two pairs of dsRNAs were prepared and purified using the RNeasy mini kit according to the manufacture’s instruction.

### Enzymatic assay in S2 cells

Vectors for exogenous expression of *HsCOX1*, *HsCOX2*, *cPges*/*p23*, and *HsPTGES3* in S2 cells were generated from *pCMV6-HsCOX1* (ORIGENE), *pCMV6-HsCOX2* (ORIGENE), LD23532 (DGRC), and *pCMV6-HsPTGES3* (ORIGENE), respectively, and cloned into the *pBRAcPA* vector. S2 cells at a density of 1 million cells in 2 ml of the culture medium were seeded on a 6-well clear flat bottom multiple well plate (Corning). For enzyme overexpression analysis, 0.5 µg of *pBRAcPA-COX1* or *pBRAcPA-COX2* and 0.5 µg of *pBRAcPA-cPges*/*p23* or *pBRAcPA*-*HsPTGES3* were transfected as described above. For RNAi experiments, 0.5 µg of *pBRAcPA-COX2* was transfected, and 10 µg of each *cPges*/*p23* dsRNA was added at the time of transfection and every 24 hours afterward. After 2 days (overexpression) or 3 days (RNAi) of incubation, 0.5 million cells in the assay medium were transferred into 24-well clear flat bottom multiple well plate (Corning) and incubated for 18 hours. Cells were incubated with selected concentrations of AA (Millipore-Sigma, #A3611) for 30 min, and then the amounts of PGE2 in the culture medium were quantified using the Prostaglandin E2 ELISA Kit (Cayman, #514010) using a VICTOR X3 luminometer (Perkin Elmer). For testing cross-reactivity of the anti-PGE2 antibody (S13A Fig), 5-hydroxyeicosatetraenoic acid (5-HETE; #34210), 9-HETE (#34400), 15-HETE (#34700), 12-hydroxyheptadecatrienoic acid (#34590), 8(9)-eicosatrienoic acid (#50351), and 11(12)-dihydroxyeicosatetraenoic acid (#51511) were obtained from Cayman Chemical.

## Data deposition

Sequence data obtained by RNA-seq are available under BioProject accession number PRJNA1126659 at the SRA (https://www.ncbi.nlm.nih.gov/sra).

## Acknowledgement

We thank Vienna *Drosophila* Resource Center, Bloomington *Drosophila* Stock Center (NIH P40 OD018537), T. Neufeld, and M.B. O’Connor for fly stocks; FlyBase (supported by NIH U41 HG000739 and U24 HG010859) for providing curated *Drosophila* genome information, S.H. Yamanaka, and M.E. Adams for sharing luminometers; Lipidomics core at the University of California, San Diego for eicosanoid determination; *Drosophila* Genomics Resource Center (NIH P40 OD010949) for cell lines; and *Drosophila* Genomics Resource Center and National Institute of Genetics for vectors and cDNA clones. This study was supported by a Pew Biomedical Scholars Award from the Pew Charitable Trusts to N.Y., a USDA NIFA Hatch project CA-R-ENT-5094-H to N.Y., an NIH Director’s New Innovator Award DP2 GM132929 to N.Y., and an NIH grant R35 GM153331 from NIGMS to N.Y.

## Author contributions

Conceptualization, N.Y.; Methodology, D.F and N.Y.; Investigation, D.F. C.N., and N.Y; Writing – Original Draft, D.F. and N.Y.; Writing – Review & Editing, D.F. and N.Y.; Supervision, N.Y.; Funding Acquisition, N.Y.

## Captions of supporting information

**S1 Fig. Phylogenetic tree of insect class A GPCRs.**

Unrooted maximum-likelihood phylogenetic tree of class A GPCRs in *Drosophila melanogaster*, *Bombyx mori*, *Apis mellifera*, *Tribolium castaneum*, *Acyrthosiphon pisum*, and *Zootermopsis nevadensis*. CG7497/PGR orthologs are conserved in all insect species analyzed. The scale bar indicates an evolutionary distance of 0.5 amino acid substitutions per site. Accession numbers of the receptors analyzed are listed in S2 Table.

**S2 Fig. Characterization of the *PGR-Gal4* strain.**

(A) Schematic diagram of the *PGR^Gal4^* allele [29]. *T2A-Gal4* is inserted in the first intron of *PGR* along with the *3xP3-GFP* marker. (B) Developmental phenotype of the *PGR^Gal4^*mutant. Most homozygous and transheterozygous *PGR* mutants died as pharate adults, indicating the loss of PGR function in *PGR^Gal4^* flies. \*\*\**p* < 0.001 (multiple comparison Chi-square test with Bonferroni correction). Numbers above the bars indicate flies analyzed in each genotype.

**S3 Fig. Expression of *PGR* during development.**

Expression patterns of *PGR* visualized by *PGR*-*Gal4*-driven expression of *UAS-mCD8::GFP*. Strong GFP signals were observed in the tracheae in all developmental stages tested. They were also observed in oenocytes in larvae, pharate adults, and newly emerged adults as indicated by arrows. Due to the *3xP3-GFP* marker in the *PGR*-*Gal4* line, background GFP signals are observed in the larval CNS, larval gut, and adult eyes as indicated by arrowheads. Detailed tissue-specific expression patterns are shown in Fig 2A. L1–L3, 1st, 2nd, and 3rd instar larvae; AE, after eclosion. Scale bars: 200 µm.

**S4 Fig. Abdominal tracheae in *PGR* mutants visualized by *UAS-mCD8::GFP*.**

(A) Pupal abdominal tracheae in heterozygous and transheterozygous *PGR* mutants visualized by *Trh-Gal4*-driven *UAS-mCD8::GFP* expression at 72 hours after puparium formation (APF). (B) Total tracheal length in the third (A3) and fourth (A4) abdominal segments in the selected area as visualized by *Trh-Gal4*-driven *UAS-mCD8::GFP* expression. Transheterozygous mutants showed defective tracheal development in both segments. n = 8–9. \*\*\**p* < 0.01 (Student’s *t*-test). (C) Pupal abdominal tracheae in heterozygous and transheterozygous *PGR* mutants visualized by *btl-Gal4*- and *dSRF-Gal4*-driven *UAS-mCD8::GFP* expression at 72 hours APF. Overexpression of *PGR* in the tracheae and TTCs partially rescued underdeveloped tracheoles in *PGR* mutants. (D) Total tracheal length in A4 in the selected area as visualized by *btl-Gal4*- and *dSRF-Gal4*-driven *UAS-mCD8::GFP* expression. Tracheal length in A3 was not measured due to excessive GFP signals in A3 driven by *dSRF-Gal4*. n = 10–11. \*\**p* < 0.1, \*\*\**p* < 0.01 (Tukey’s honestly significant difference test). Scale bars: 200 µm.

**S5 Fig. Phylogenetic tree of heme peroxidases in *H. sapiens* and insects.**

Unrooted maximum-likelihood phylogenetic tree of heme peroxidases in *Homo sapiens*, *Drosophila melanogaster*, *Bombyx mori*, *Apis mellifera*, *Tribolium castaneum*, *Acyrthosiphon pisum*, and *Zootermopsis nevadensis*. Branches are color-coded for different species.

Cyclooxygenases (PGG/H synthases) in *H. sapiens* are highlighted. The scale bar indicates an evolutionary distance of 0.5 amino acid substitutions per site. Accession numbers of the enzymes analyzed are listed in S3 Table.

**S6 Fig. Phylogenetic tree of glutathione S-transferases in *H. sapiens* and insects.**

Unrooted maximum-likelihood phylogenetic tree of glutathione S-transferases in *Homo sapiens*, *Drosophila melanogaster*, *Bombyx mori*, *Apis mellifera*, *Tribolium castaneum*, *Acyrthosiphon pisum*, and *Zootermopsis nevadensis*. Branches are color-coded for different species. Clades that include PGD synthase and PGE synthase 1 in *H. sapiens* are highlighted. The scale bar indicates an evolutionary distance of 0.5 amino acid substitutions per site. Accession numbers of the enzymes analyzed are listed in S4 Table.

**S7 Fig. Phylogenetic tree of glutaredoxins in *H. sapiens* and insects.**

Unrooted maximum-likelihood phylogenetic tree of glutaredoxin domain-containing proteins in *Homo sapiens*, *Drosophila melanogaster*, *Bombyx mori*, *Apis mellifera*, *Tribolium castaneum*, *Acyrthosiphon pisum*, and *Zootermopsis nevadensis*. Branches are color-coded for different species. The clade that includes PGE synthase 2 in *H. sapiens* is highlighted. The scale bar indicates an evolutionary distance of 0.5 amino acid substitutions per site. Accession numbers of the enzymes analyzed are listed in S5 Table.

**S8 Fig. Phylogenetic tree of carbonyl reductases in *H. sapiens* and insects.**

Unrooted maximum-likelihood phylogenetic tree of carbonyl reductases in *Homo sapiens*, *Drosophila melanogaster*, *Bombyx mori*, *Apis mellifera*, *Tribolium castaneum*, *Acyrthosiphon pisum*, and *Zootermopsis nevadensis*. Branches are color-coded for different species. The clade that includes carbonyl reductase 1 (PGF synthase) in *H. sapiens* is highlighted. The scale bar indicates an evolutionary distance of 0.5 amino acid substitutions per site. Accession numbers of the enzymes analyzed are listed in S6 Table.

**S9 Fig. Phylogenetic tree of aldo-keto reductases in *H. sapiens* and insects.**

Unrooted maximum-likelihood phylogenetic tree of aldo-keto reductases in *Homo sapiens*, *Drosophila melanogaster*, *Bombyx mori*, *Apis mellifera*, *Tribolium castaneum*, *Acyrthosiphon pisum*, and *Zootermopsis nevadensis*. Branches are color-coded for different species. The clade that includes aldo-keto reductase (PGF synthase 2) in *H. sapiens* is highlighted. The scale bar indicates an evolutionary distance of 0.5 amino acid substitutions per site. Accession numbers of the enzymes analyzed are listed in S7 Table.

**S10 Fig. Phylogenetic tree of peroxiredoxins in *H. sapiens* and insects.**

Unrooted maximum-likelihood phylogenetic tree of peroxiredoxins in *Homo sapiens*, *Drosophila melanogaster*, *Bombyx mori*, *Apis mellifera*, *Tribolium castaneum*, *Acyrthosiphon pisum*, and *Zootermopsis nevadensis*. Branches are color-coded for different species. The clade that includes peroxiredoxin (PGF synthase) in *H. sapiens* is highlighted. The scale bar indicates an evolutionary distance of 0.5 amino acid substitutions per site. Accession numbers of the enzymes analyzed are listed in S8 Table.

**S11 Fig. Phylogenetic tree of cytochrome P450 enzymes in *H. sapiens* and insects.**

Unrooted maximum-likelihood phylogenetic tree of cytochrome P450 enzymes in *Homo sapiens*, *Drosophila melanogaster*, *Bombyx mori*, *Apis mellifera*, *Tribolium castaneum*, *Acyrthosiphon pisum*, and *Zootermopsis nevadensis*. Branches are color-coded for different species. The four major CYP clans in insects are highlighted. There are no orthologous enzymes of Cyp5A1 or Cyp8A1 in insects. The scale bar indicates an evolutionary distance of 0.5 amino acid substitutions per site. Accession numbers of the enzymes analyzed are listed in S9 Table.

**S12 Fig. Mutagenesis of PGD/PGE synthase orthologs.**

Mutagenesis was conducted by CRISPR-Cas9-based homologous recombination to insert the 3xP3-RFP sequence into each target site. Two single guide RNAs (sgRNAs) were designed for each target to delete entire coding sequences shown in orange. Genomic sequences of the mutants are provided in S2–S6 Documents.

**S13 Fig. ELISA specificity and knockdown efficiency regarding the enzymatic conversion assay in *Drosophila* S2 cells.**

(A) Cross-reactivity of the anti-PGE2 antibody used in the PGE2 ELISA system. Serial dilutions of AA and various eicosanoids previously detected in the *Drosophila* hemolymph after AA injection [22] were tested for their potential cross-reactivity. AA and PGF2α have 0.017% and 0.25% cross-reactivity, respectively. HETE, hydroxyeicosatetraenoic acid; HHTrE, hydroxyheptadecatrienoic acid; EET, eicosatrienoic acid; DiHET, dihydroxyeicosatetraenoic acid. (B) Relative expression levels of *cPges*/*p23* in S2 cells treated with dsRNA for 3 days*. cPges*/*p23* mRNA levels were downregulated in *cPges*/*p23* RNAi cells as compared to the negative control (*EGFP* RNAi). Expression levels are normalized by the levels of a reference gene, *rp49*, in the same cDNA samples. n = 3. \**p* < 0.05, ** < 0.01 (Dunnett’s test *vs EGFP* RNAi).

**S14 Fig. Hypoxia response and tracheogenesis in *PGR* mutants.**

(A) Relative expression levels of *branchless* (*bnl*) in *PGR* mutants from 0 to 96 hours after puparium formation (APF). Insects pupated about 12 hours APF. Homozygous mutant pupae did not show significantly higher expression of *bnl* until 96 hours APF. (B, C) Relative expression levels of hypoxia response genes (*Ldh* and *Hph*) and *bnl* in adult *PGR* mutants rescued by high oxygen supply during pupa-adult development. Eclosed flies were transferred to the normal oxygen condition within 2 hours after eclosion and kept there for 30 min (B) or 24 hours (C) before RNA extraction. Rescued *PGR* mutant flies express high levels of *Ldh* and *Hph* immediately after eclosion, which decreases within 24 hours. In contrast, *bnl* continues to be highly expressed in *PGR* mutant flies 24 hours after eclosion. Expression levels are normalized by the levels of a reference gene, *rp49*, in the same cDNA samples and shown as relative to *PGR^A1/+^* at 0 hours APF. n = 3–4. \**p* < 0.05, \*\**p* < 0.01, \*\*\**p* < 0.001 (Student’s *t-*test *vs PGR^A1/+^*). (D) Adult tracheal development in *PGR* heterozygous and transheterozygous mutants raised under 40% oxygen during pupa-adult development. Abdominal tracheae in adult males were visualized by *btl-Gal4*-driven *UAS-mCD8::GFP* expression within 2 hours or 24 hours after eclosion. In newly emerged *PGR* transheterozygous mutants, the adult tracheal system was underdeveloped, which recovered to the level of the control heterozygous mutants within 24 hours after eclosion. Scale bar: 200 µm.

**S15 Fig. Representative antimicrobial peptide and reproduction-related genes downregulated in *PGR* mutants.**

Expression of antimicrobial peptide genes (A) and reproduction-related genes (B) in *w^1118^* control and the *PGR* mutant based on RNA-seq data. Expression levels of *Cecropin A1* and *A2* (*CecA1* and *CecA2*), *Defensin* (*Def*), and *Drosomycin* (*Drs*) in the *PGR* mutant were significantly lower than control at 0 hours after puparium formation (APF). Expression of *Sex peptide receptor* (*SPR*), *Yolk protein 1* (*Yp1*), and *Yp2* in the *PGR* mutant was significantly downregulated at 96 hours APF. Expression levels are shown as reads per kilobase of transcript per million mapped reads (RPKM). \**p* < 0.05, \*\*\**p* < 0.001 (Student’s *t-*test).

**S1 Table. GPCRs in the phylogenetic tree, related to Figs 1A and 1B**.

**S2 Table. GPCRs in the phylogenetic tree, related to S1 Fig.**

**S3 Table. Heme peroxidases in the phylogenetic tree, related to S4 Fig.**

**S4 Table. Glutathione S-transferases in the phylogenetic tree, related to S5 Fig.**

**S5 Table. Glutaredoxins in the phylogenetic tree, related to S6 Fig.**

**S6 Table. Carbonyl reductases in the phylogenetic tree, related to S7 Fig.**

**S7 Table. Aldo-keto reductases in the phylogenetic tree, related to S8 Fig.**

**S8 Table. Peroxiredoxins in the phylogenetic tree, related to S9 Fig.**

**S9 Table. Cytochrome P450 enzymes in the phylogenetic tree, related to S10 Fig.**

**S10 Table. Top affected gene functions in the *PGR* mutant.**

**S11 Table: *Drosophila* strains.**

**S12 Table: Plasmid constructs.**

**S13 Table: Primers and oligonucleotides.**

