## Supporting Information for "Functional characterization of eicosanoid signaling in *Drosophila* development"

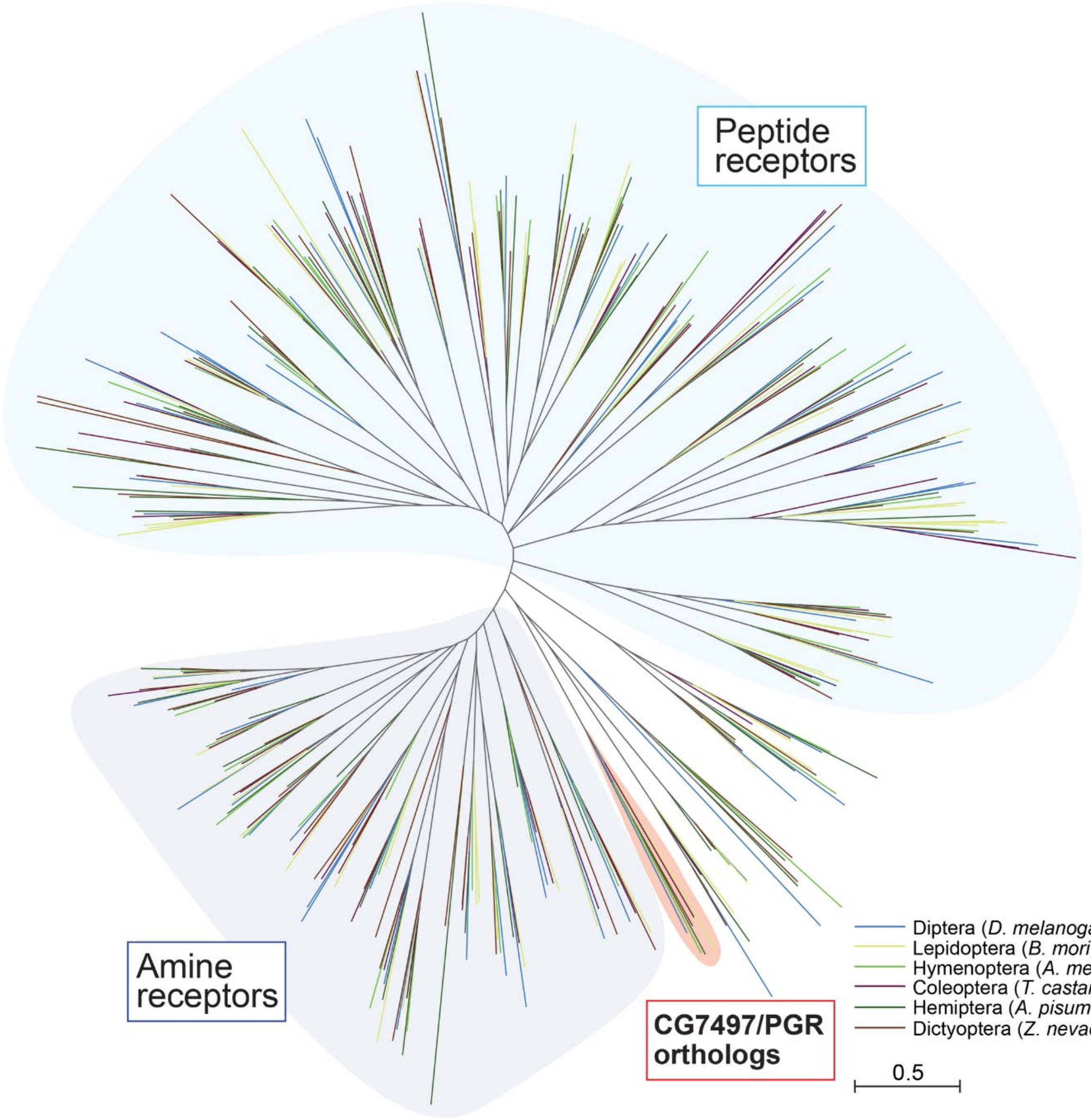


**S1 Fig. Phylogenetic tree of insect class A GPCRs.**

Unrooted maximum-likelihood phylogenetic tree of class A GPCRs in *Drosophila melanogaster*, *Bombyx mori*, *Apis mellifera*, *Tribolium castaneum*, *Acyrthosiphon pisum*, and *Zootermopsis nevadensis*. CG7497/PGR orthologs are conserved in all insect species analyzed. The scale bar indicates an evolutionary distance of 0.5 amino acid substitutions per site. Accession numbers of the receptors analyzed are listed in S2 Table.

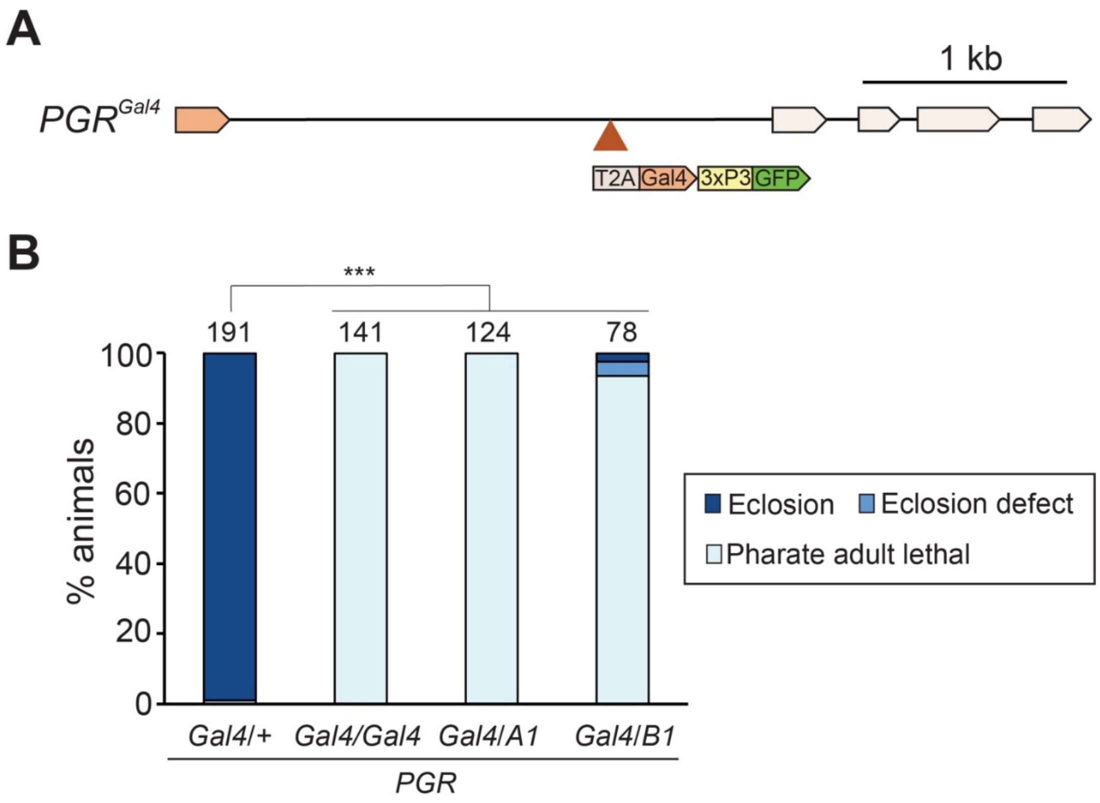


**S2 Fig. Characterization of the *PGR-Gal4* strain.**

(**A**) Schematic diagram of the $PGR^{Gal4}$ allele [29]. *T2A-Gal4* is inserted in the first intron of *PGR* along with the *3xP3-GFP* marker. (**B**) Developmental phenotype of the $PGR^{Gal4}$ mutant. Most homozygous and transheterozygous *PGR* mutants died as pharate adults, indicating the loss of PGR function in $PGR^{Gal4}$ flies. ***$p < 0.001$ (multiple comparison Chi-square test with Bonferroni correction). Numbers above the bars indicate flies analyzed in each genotype.

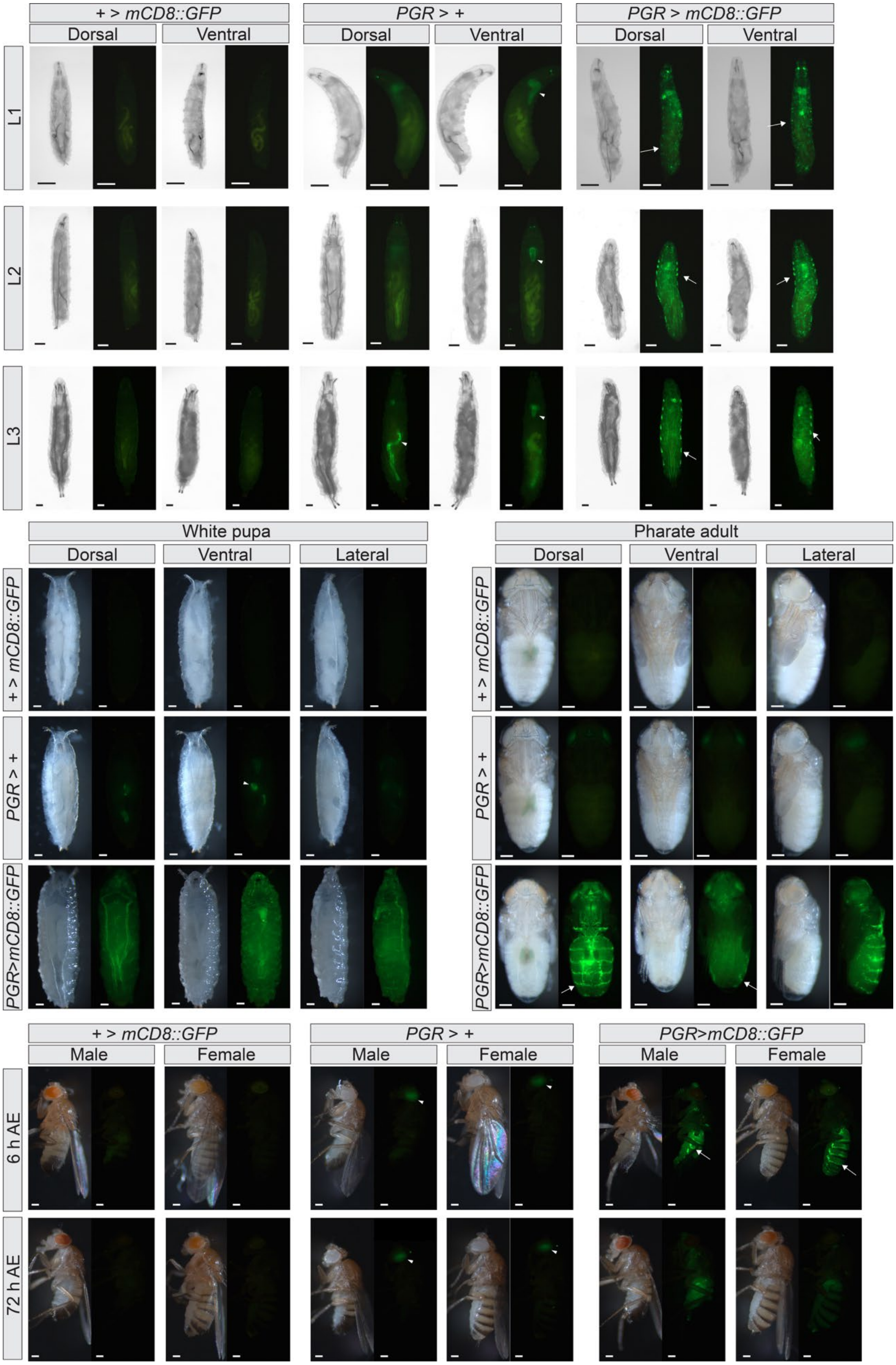
+ > mCD8::GFP
PGR > +
PGR > mCD8::GFP
Dorsal
Ventral
L1
L2
L3
White pupa
Pharate adult
Lateral
PGR>mCD8::GFP
Male
Female
6 h AE
72 h AE

**S3 Fig. Expression of *PGR* during development.**

Expression patterns of *PGR* visualized by *PGR-Gal4*-driven expression of *UAS-mCD8::GFP*. Strong GFP signals were observed in the tracheae in all developmental stages tested. They were also observed in oenocytes in larvae, pharate adults, and newly emerged adults as indicated by arrows. Due to the *3xP3-GFP* marker in the *PGR-Gal4* line, background GFP signals are observed in the larval CNS, larval gut, and adult eyes as indicated by arrowheads. Detailed tissue-specific expression patterns are shown in Fig 2A. L1–L3, 1st, 2nd, and 3rd instar larvae; AE, after eclosion. Scale bars: 200 µm.

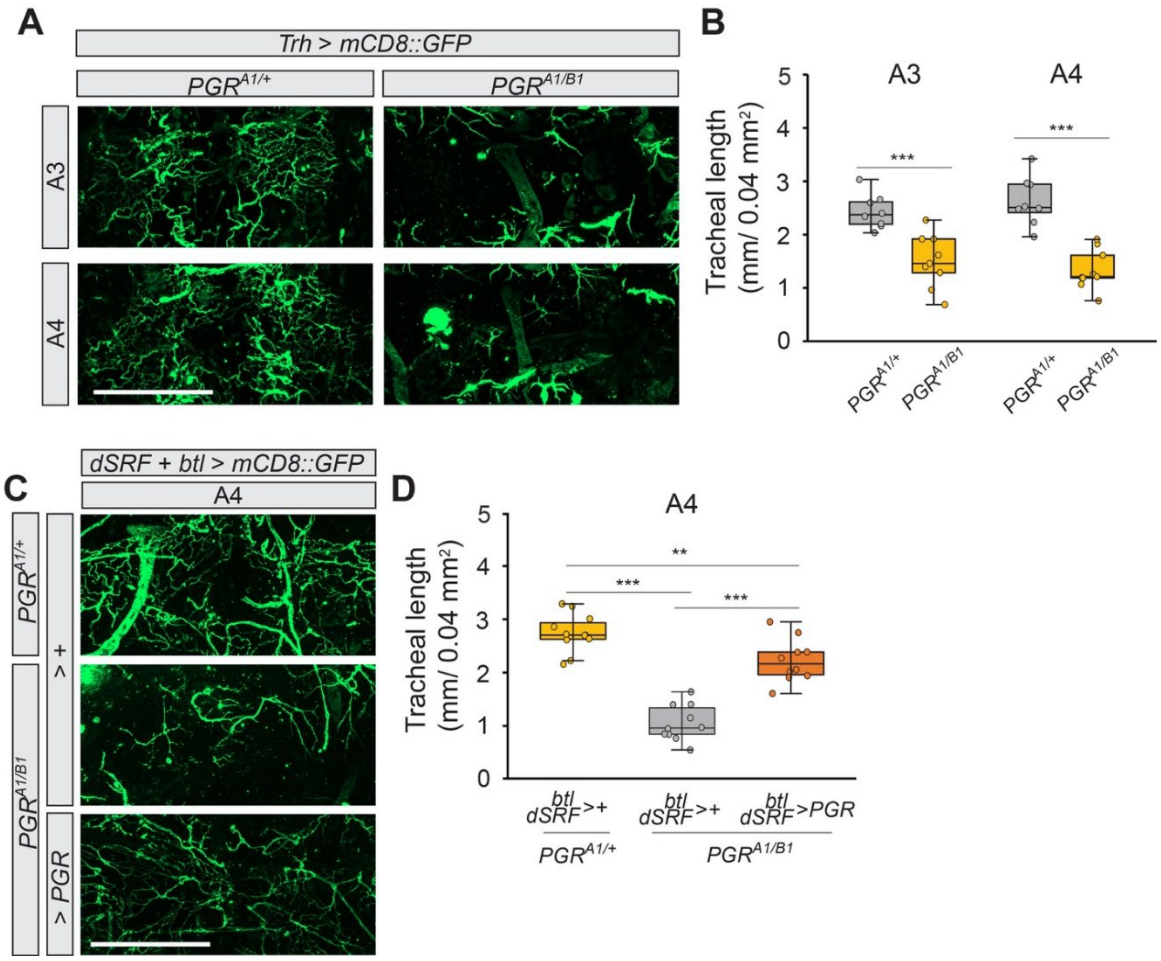


**S4 Fig. Abdominal tracheae in *PGR* mutants visualized by *UAS-mCD8::GFP*.**

(**A**) Pupal abdominal tracheae in heterozygous and transheterozygous *PGR* mutants visualized by *Trh-Gal4*-driven *UAS-mCD8::GFP* expression at 72 hours after puparium formation (APF). (**B**) Total tracheal length in the third (A3) and fourth (A4) abdominal segments in the selected area as visualized by *Trh-Gal4*-driven *UAS-mCD8::GFP* expression. Transheterozygous mutants showed defective tracheal development in both segments. n = 8–9. ***$p < 0.01$ (Student's *t*-test). (**C**) Pupal abdominal tracheae in heterozygous and transheterozygous *PGR* mutants visualized by *btl-Gal4*- and *dSRF-Gal4*-driven *UAS-mCD8::GFP* expression at 72 hours APF. Overexpression of *PGR* in the tracheae and TTCs partially rescued underdeveloped tracheoles in *PGR* mutants. (**D**) Total tracheal length in A4 in the selected area as visualized by *btl-Gal4*- and *dSRF-Gal4*-driven *UAS-mCD8::GFP* expression. Tracheal length in A3 was not measured due to excessive GFP signals in A3 driven by *dSRF-Gal4*. n = 10–11. **$p < 0.1$, ***$p < 0.01$ (Tukey's honestly significant difference test). Scale bars: 200 µm.

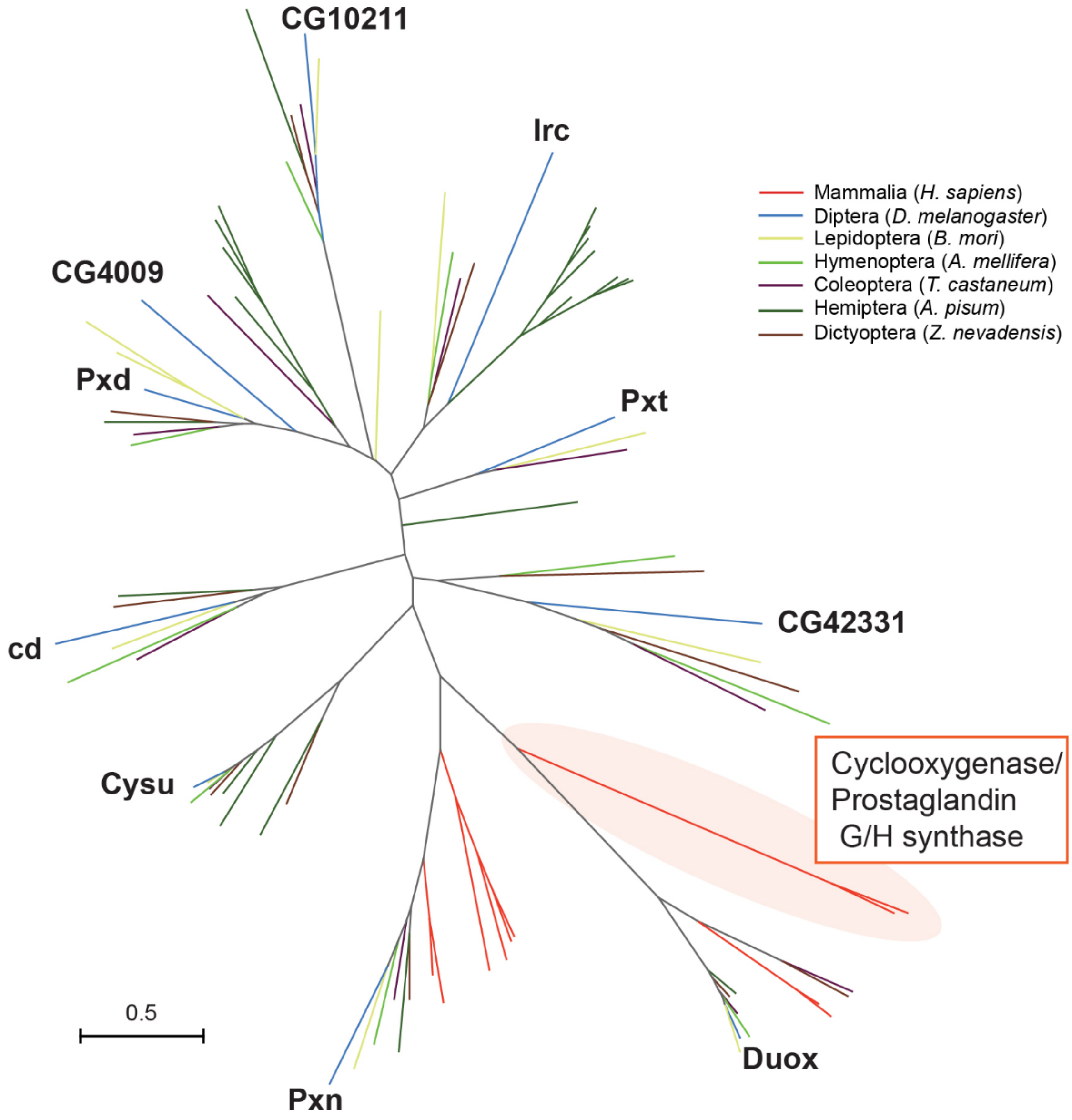


**S5 Fig. Phylogenetic tree of heme peroxidases in *H. sapiens* and insects.**

Unrooted maximum-likelihood phylogenetic tree of heme peroxidases in *Homo sapiens*, *Drosophila melanogaster*, *Bombyx mori*, *Apis mellifera*, *Tribolium castaneum*, *Acyrthosiphon pisum*, and *Zootermopsis nevadensis*. Branches are color-coded for different species. Cyclooxygenases (PGG/H synthases) in *H. sapiens* are highlighted. The scale bar indicates an evolutionary distance of 0.5 amino acid substitutions per site. Accession numbers of the enzymes analyzed are listed in S3 Table.

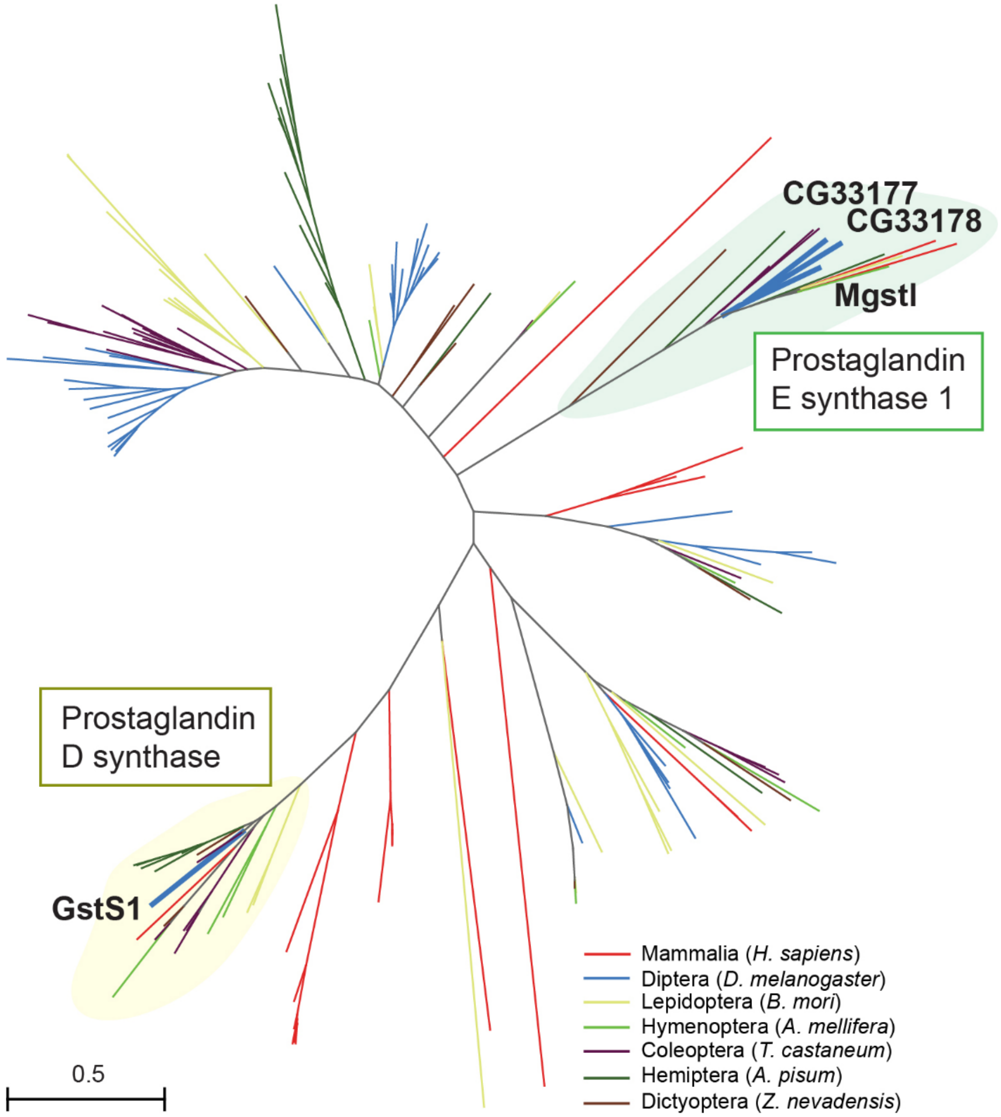


**S6 Fig. Phylogenetic tree of glutathione S-transferases in *H. sapiens* and insects.**

Unrooted maximum-likelihood phylogenetic tree of glutathione S-transferases in *Homo sapiens*, *Drosophila melanogaster*, *Bombyx mori*, *Apis mellifera*, *Tribolium castaneum*, *Acyrthosiphon pisum*, and *Zootermopsis nevadensis*. Branches are color-coded for different species. Clades that include PGD synthase and PGE synthase 1 in *H. sapiens* are highlighted. The scale bar indicates an evolutionary distance of 0.5 amino acid substitutions per site. Accession numbers of the enzymes analyzed are listed in S4 Table.

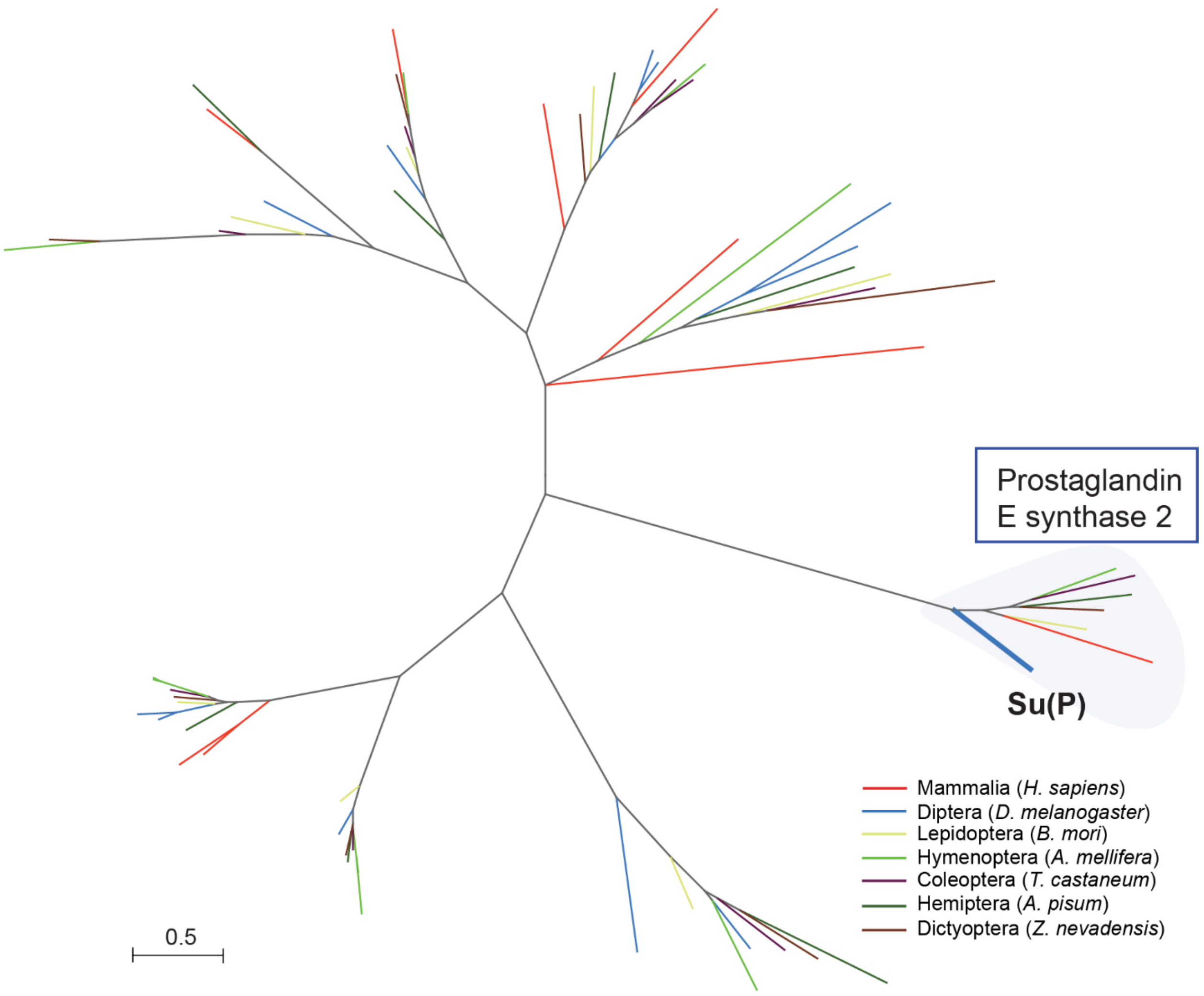


**S7 Fig. Phylogenetic tree of glutaredoxins in *H. sapiens* and insects.**

Unrooted maximum-likelihood phylogenetic tree of glutaredoxin domain-containing proteins in *Homo sapiens*, *Drosophila melanogaster*, *Bombyx mori*, *Apis mellifera*, *Tribolium castaneum*, *Acyrthosiphon pisum*, and *Zootermopsis nevadensis*. Branches are color-coded for different species. The clade that includes PGE synthase 2 in *H. sapiens* is highlighted. The scale bar indicates an evolutionary distance of 0.5 amino acid substitutions per site. Accession numbers of the enzymes analyzed are listed in S5 Table.

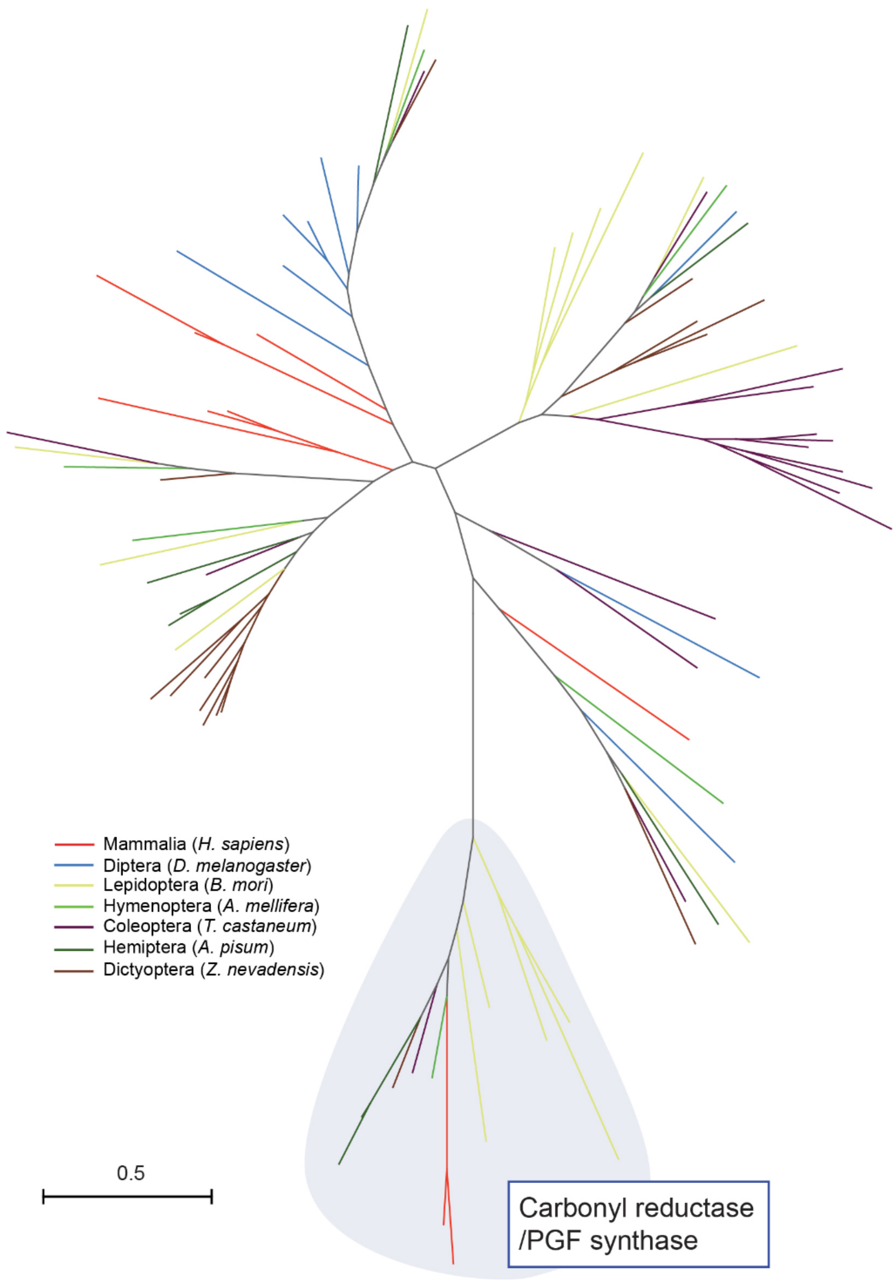


**S8 Fig. Phylogenetic tree of carbonyl reductases in *H. sapiens* and insects.**

Unrooted maximum-likelihood phylogenetic tree of carbonyl reductases in *Homo sapiens*, *Drosophila melanogaster*, *Bombyx mori*, *Apis mellifera*, *Tribolium castaneum*, *Acyrthosiphon pisum*, and *Zootermopsis nevadensis*. Branches are color-coded for different species. The clade that includes carbonyl reductase 1 (PGF synthase) in *H. sapiens* is highlighted. The scale bar indicates an evolutionary distance of 0.5 amino acid substitutions per site. Accession numbers of the enzymes analyzed are listed in S6 Table.

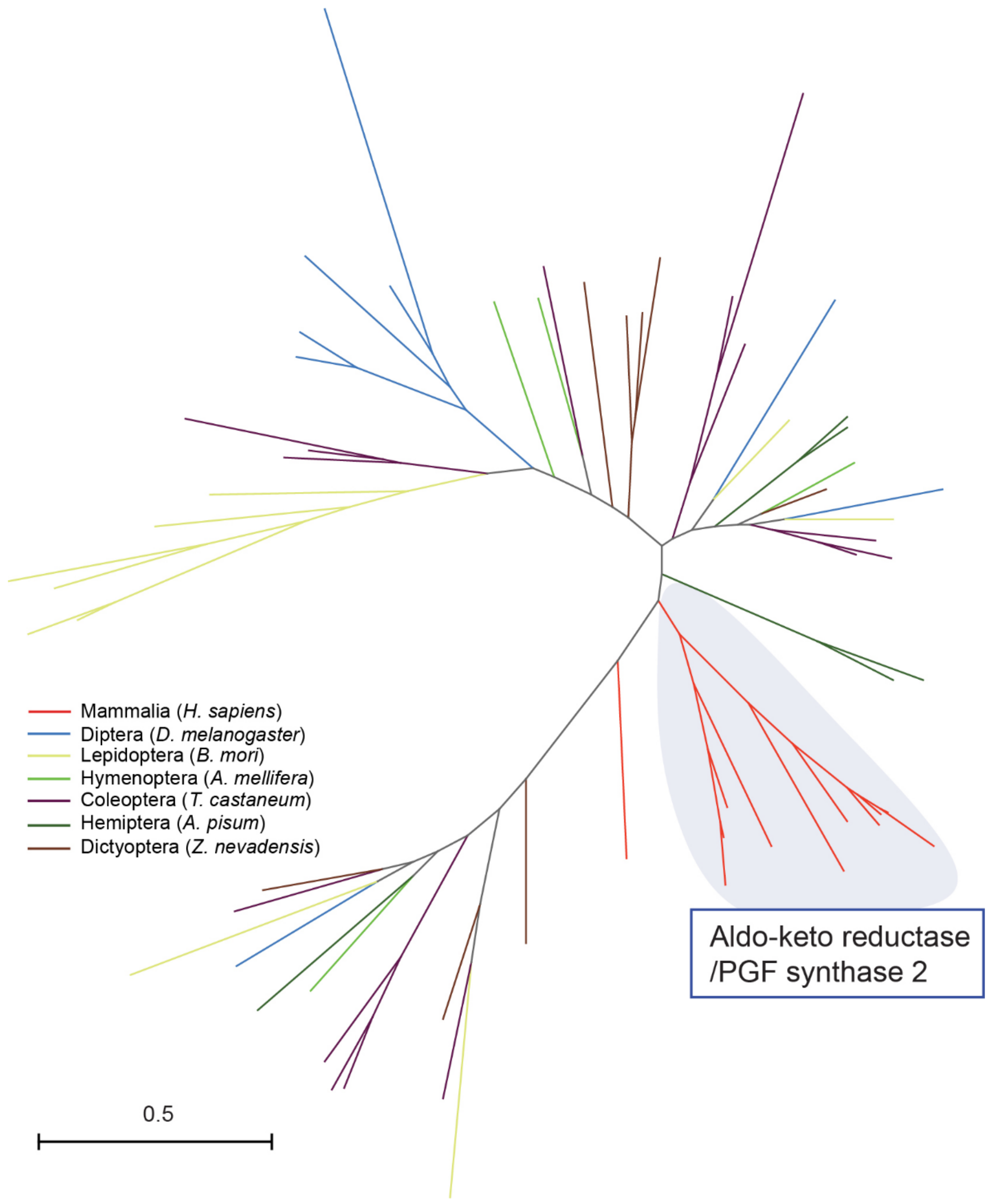


**S9 Fig. Phylogenetic tree of aldo-keto reductases in *H. sapiens* and insects.**

Unrooted maximum-likelihood phylogenetic tree of aldo-keto reductases in *Homo sapiens*, *Drosophila melanogaster*, *Bombyx mori*, *Apis mellifera*, *Tribolium castaneum*, *Acyrthosiphon pisum*, and *Zootermopsis nevadensis*. Branches are color-coded for different species. The clade that includes aldo-keto reductase (PGF synthase 2) in *H. sapiens* is highlighted. The scale bar indicates an evolutionary distance of 0.5 amino acid substitutions per site. Accession numbers of the enzymes analyzed are listed in S7 Table.

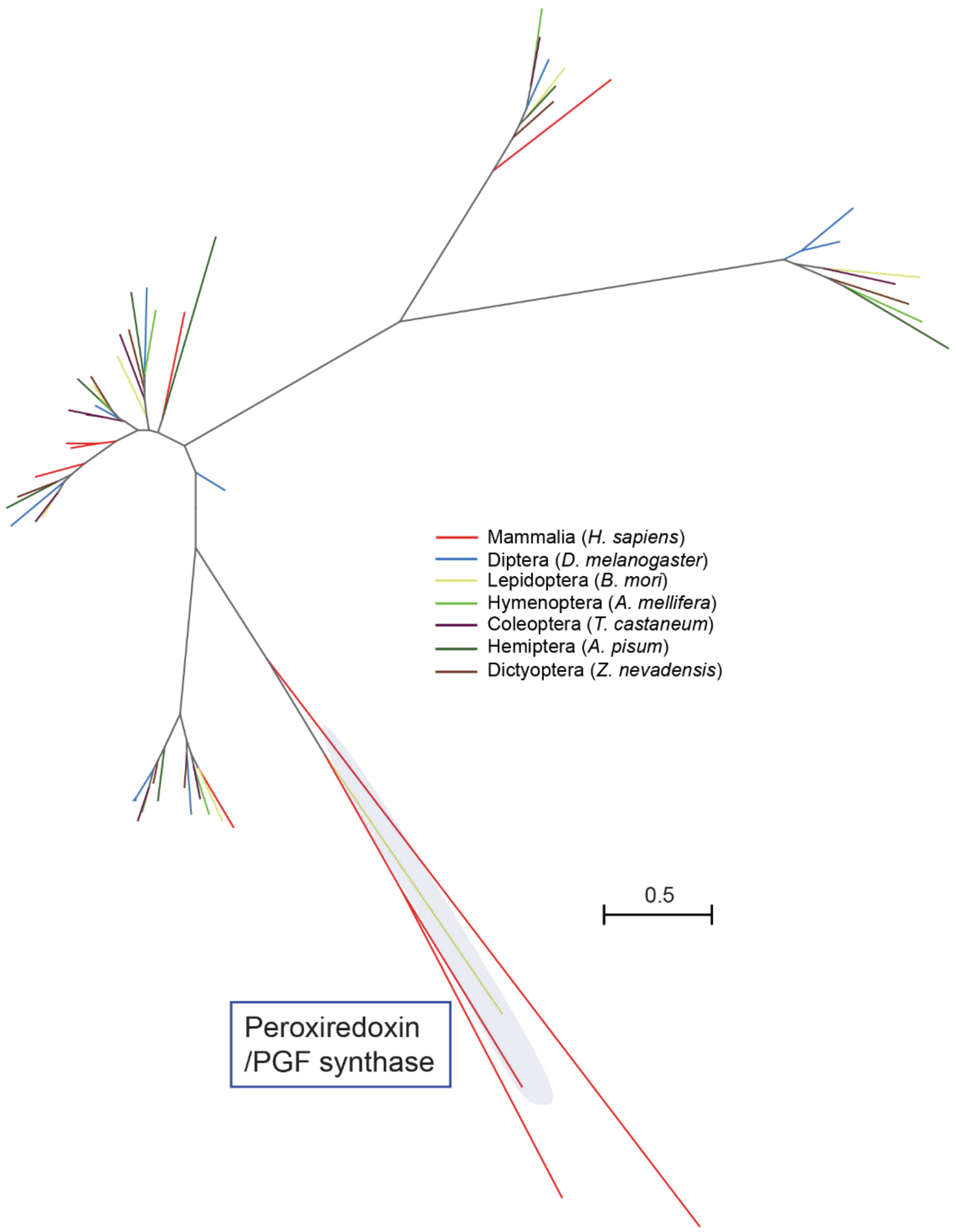


**S10 Fig. Phylogenetic tree of peroxiredoxins in *H. sapiens* and insects.**

Unrooted maximum-likelihood phylogenetic tree of peroxiredoxins in *Homo sapiens*, *Drosophila melanogaster*, *Bombyx mori*, *Apis mellifera*, *Tribolium castaneum*, *Acyrthosiphon pisum*, and *Zootermopsis nevadensis*. Branches are color-coded for different species. The clade that includes peroxiredoxin (PGF synthase) in *H. sapiens* is highlighted. The scale bar indicates an evolutionary distance of 0.5 amino acid substitutions per site. Accession numbers of the enzymes analyzed are listed in S8 Table.

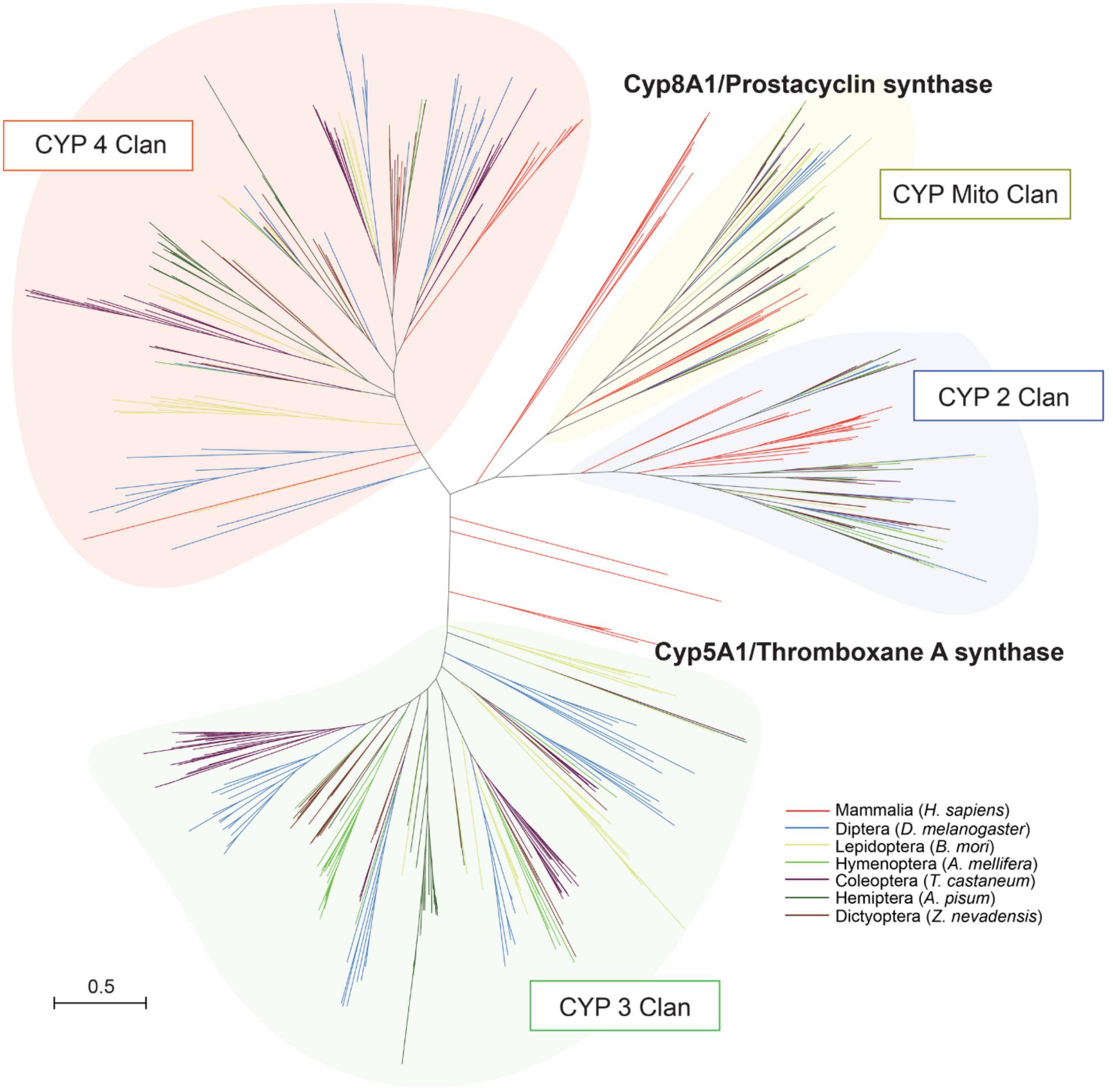


**S11 Fig. Phylogenetic tree of cytochrome P450 enzymes in *H. sapiens* and insects.**
Unrooted maximum-likelihood phylogenetic tree of cytochrome P450 enzymes in *Homo sapiens*, *Drosophila melanogaster*, *Bombyx mori*, *Apis mellifera*, *Tribolium castaneum*, *Acyrthosiphon pisum*, and *Zootermopsis nevadensis*. Branches are color-coded for different species. The four major CYP clans in insects are highlighted. There are no orthologous enzymes of Cyp5A1 or Cyp8A1 in insects. The scale bar indicates an evolutionary distance of 0.5 amino acid substitutions per site. Accession numbers of the enzymes analyzed are listed in S9 Table.

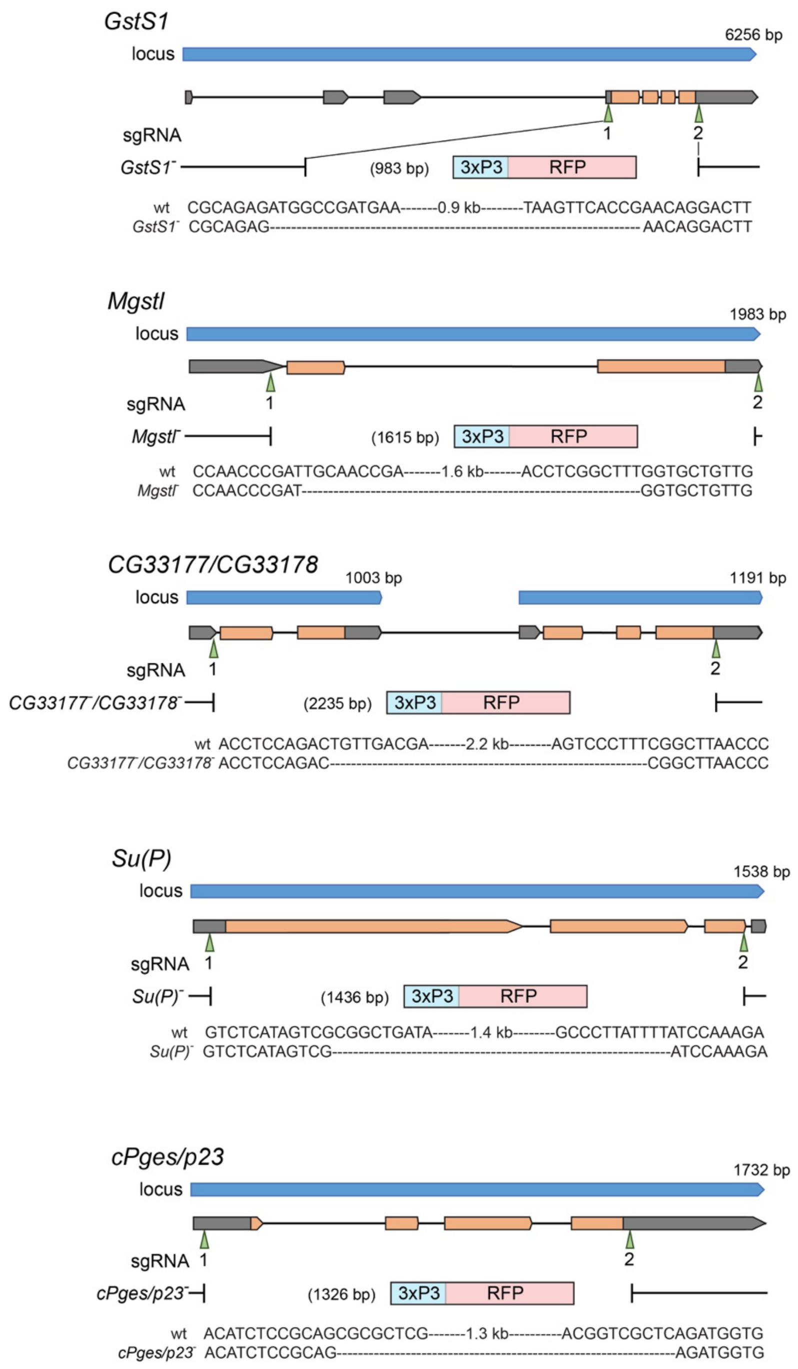


**S12 Fig. Mutagenesis of PGD/PGE synthase orthologs.**

Mutagenesis was conducted by CRISPR-Cas9-based homologous recombination to insert the 3xP3-RFP sequence into each target site. Two single guide RNAs (sgRNAs) were designed for each target to delete entire coding sequences shown in orange. Genomic sequences of the mutants are provided in S2–S6 Documents.

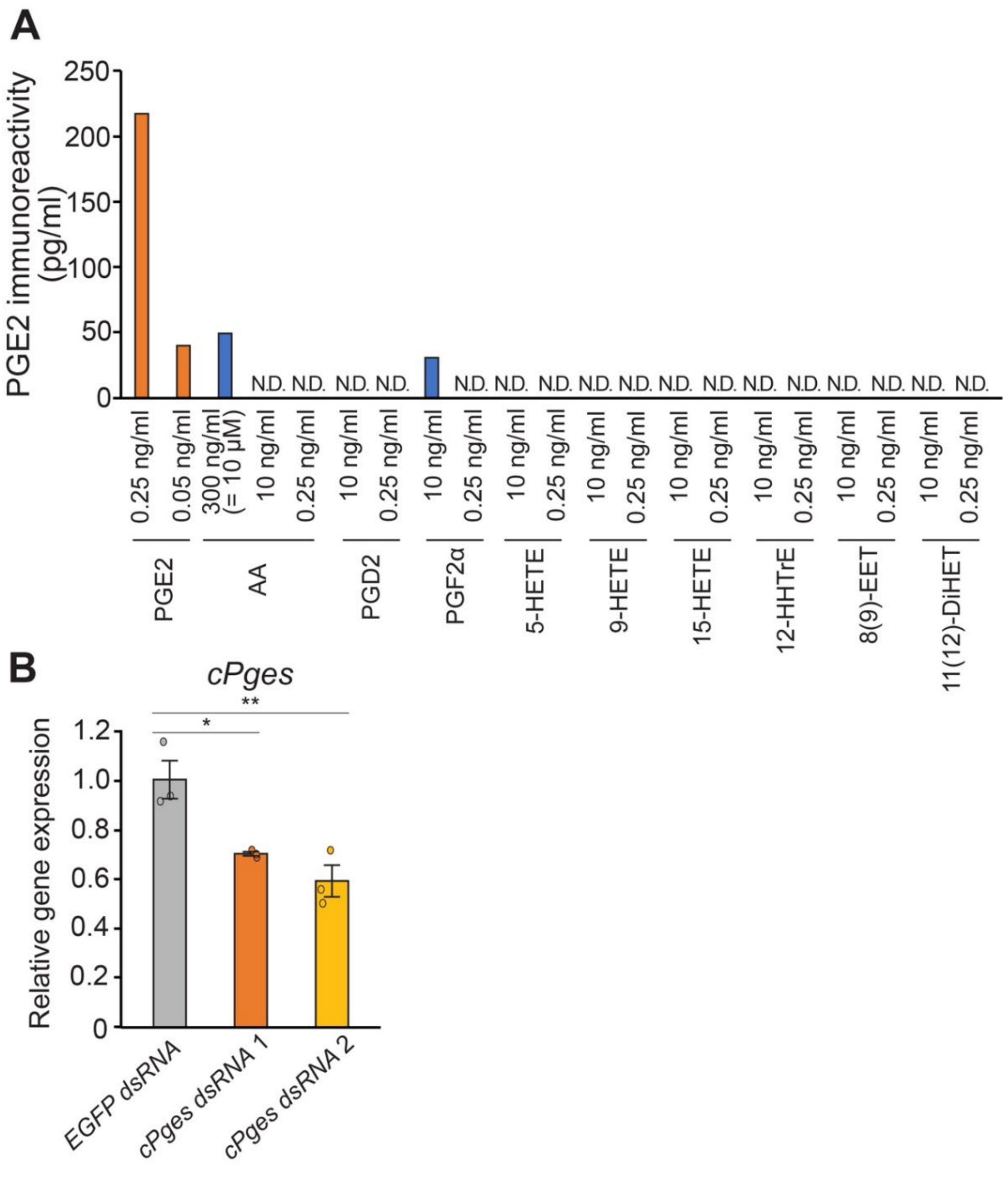


**S13 Fig. ELISA specificity and knockdown efficiency regarding the enzymatic conversion assay in *Drosophila* S2 cells.**

(**A**) Cross-reactivity of the anti-PGE2 antibody used in the PGE2 ELISA system. Serial dilutions of AA and various eicosanoids previously detected in the *Drosophila* hemolymph after AA injection [22] were tested for their potential cross-reactivity. AA and PGF2α have 0.017% and 0.25% cross-reactivity, respectively. HETE, hydroxyeicosatetraenoic acid; HHTrE, hydroxyheptadecatrienoic acid; EET, eicosatrienoic acid; DiHET, dihydroxyeicosatetraenoic acid. (**B**) Relative expression levels of *cPges*/*p23* in S2 cells treated with dsRNA for 3 days.

*cPges*/*p23* mRNA levels were downregulated in *cPges*/*p23* RNAi cells as compared to the negative control (*EGFP* RNAi). Expression levels are normalized by the levels of a reference gene, *rp49*, in the same cDNA samples. n = 3. *$p < 0.05$, ** < 0.01 (Dunnett's test *vs EGFP* RNAi).

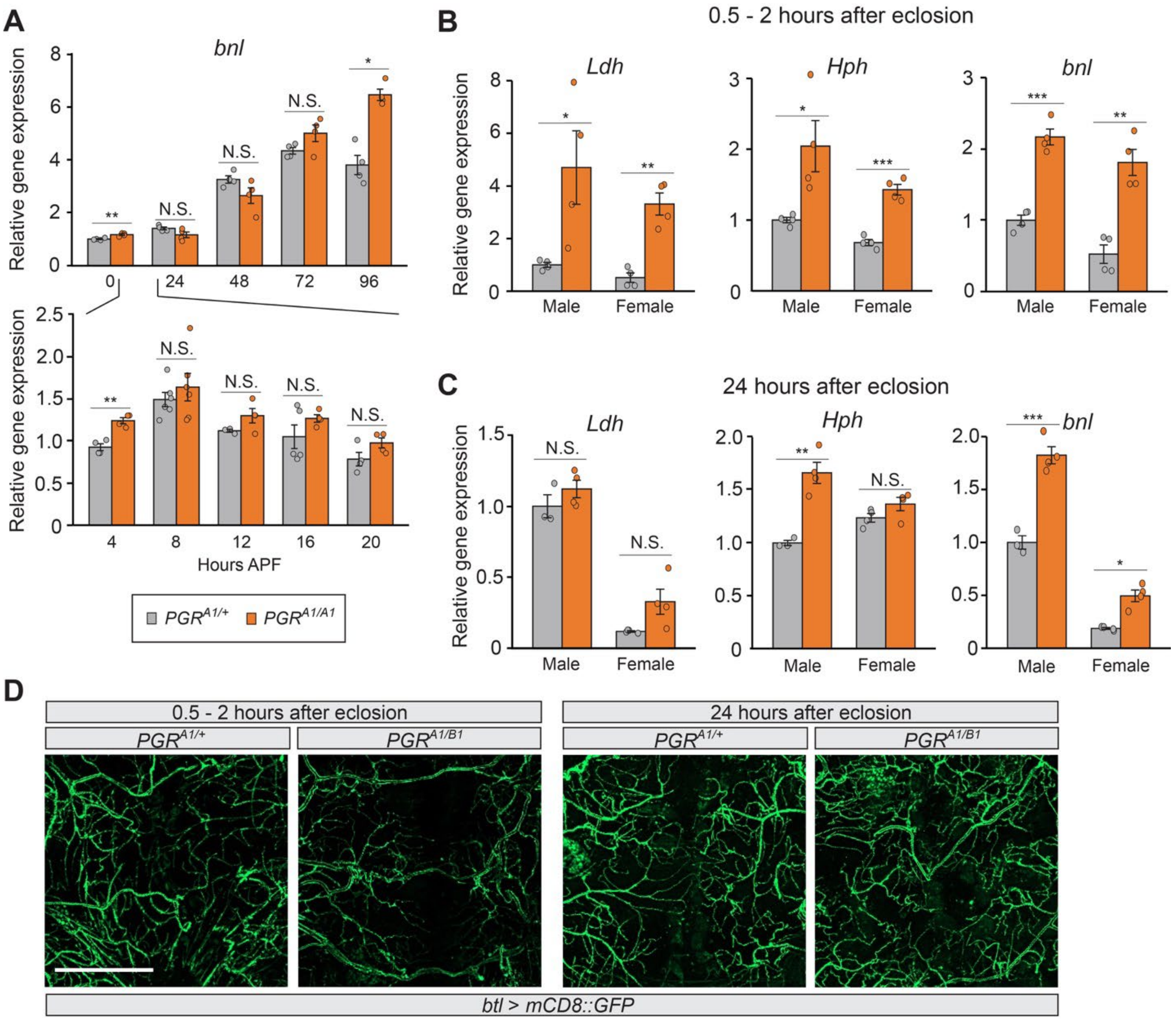


**S14 Fig. Hypoxia response and tracheogenesis in *PGR* mutants.**

(**A**) Relative expression levels of *branchless* (*bnl*) in *PGR* mutants from 0 to 96 hours after puparium formation (APF). Insects pupated about 12 hours APF. Homozygous mutant pupae did not show significantly higher expression of *bnl* until 96 hours APF. (**B, C**) Relative expression levels of hypoxia response genes (*Ldh* and *Hph*) and *bnl* in adult *PGR* mutants rescued by high oxygen supply during pupa-adult development. Eclosed flies were transferred to the normal oxygen condition within 2 hours after eclosion and kept there for 30 min (B) or 24 hours (C) before RNA extraction. Rescued *PGR* mutant flies express high levels of *Ldh* and *Hph* immediately after eclosion, which decreases within 24 hours. In contrast, *bnl* continues to be highly expressed in *PGR* mutant flies 24 hours after eclosion. Expression levels are normalized by the levels of a reference gene, *rp49*, in the same cDNA samples and shown as relative to *PGR*$^{A1/+}$ at 0 hours APF. n = 3–4. *$p$ < 0.05, **$p$ < 0.01, ***$p$ < 0.001 (Student's *t*-test *vs* *PGR*$^{A1/+}$). (**D**) Adult tracheal development in *PGR* heterozygous and transheterozygous mutants raised under 40% oxygen during pupa-adult development. Abdominal tracheae in adult males were visualized by *btl-Gal4*-driven *UAS-mCD8::GFP* expression within 2 hours or 24 hours after eclosion. In newly emerged *PGR* transheterozygous mutants, the adult tracheal system was underdeveloped, which recovered to the level of the control heterozygous mutants within 24 hours after eclosion. Scale bar: 200 µm.

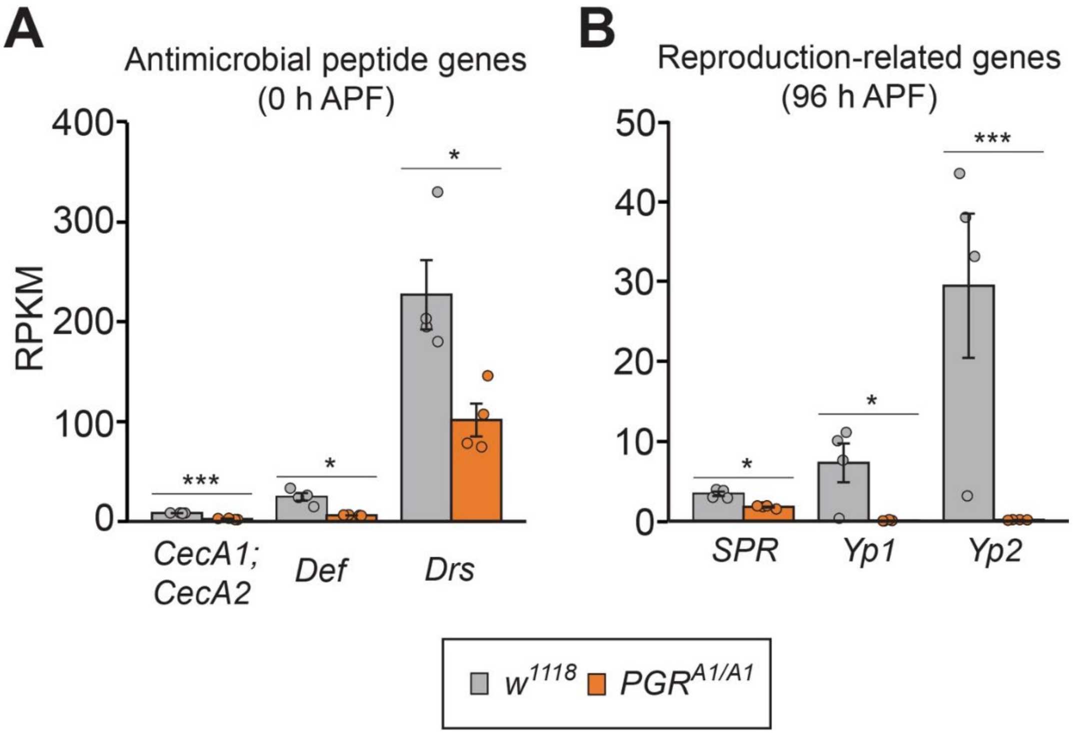


**S15 Fig. Representative antimicrobial peptide and reproduction-related genes downregulated in *PGR* mutants.**

Expression of antimicrobial peptide genes (A) and reproduction-related genes (B) in $w^{1118}$ control and the *PGR* mutant based on RNA-seq data. Expression levels of *Cecropin A1* and *A2* (*CecA1* and *CecA2*), *Defensin* (*Def*), and *Drosomycin* (*Drs*) in the *PGR* mutant were significantly lower than control at 0 hours after puparium formation (APF). Expression of *Sex peptide receptor* (*SPR*), *Yolk protein 1* (*Yp1*), and *Yp2* in the *PGR* mutant was significantly downregulated at 96 hours APF. Expression levels are shown as reads per kilobase of transcript per million mapped reads (RPKM). *$p < 0.05$, ***$p < 0.001$ (Student's *t*-test).

**S11 Table: *Drosophila* strains.**

| Organisms: Strains | Source | Identifier |
|---|---|---|
| *D. melanogaster*: $w^{1118}$ | Bloomington *Drosophila* Stock Center | BDSC: 5905 |
| *D. melanogaster*: *tubP-Gal4* (Whole body) | Bloomington *Drosophila* Stock Center | BDSC: 5138 |
| *D. melanogaster*: *PGR-Gal4* | Bloomington *Drosophila* Stock Center | BDSC: 86488 |
| *D. melanogaster*: *btl-Gal4* (Trachea) | Bloomington *Drosophila* Stock Center | BDSC: 78328 |
| *D. melanogaster*: *dSRF-Gal4* (Terminal tracheal cells) | Bloomington *Drosophila* Stock Center | BDSC: 25753 |
| *D. melanogaster*: *Trh-Gal4* (Trachea) | Bloomington *Drosophila* Stock Center | BDSC: 47463 |
| *D. melanogaster*: *elav-Gal4* (Central nervous system) | Gift from Michael B. O'Connor (University of Minnesota) | N/A |
| *D. melanogaster*: *nSyb-Gal4* (Neuron) | Bloomington *Drosophila* Stock Center | BDSC: 51941 |
| *D. melanogaster*: *Desat1-Gal4* (Oenocytes) | Bloomington *Drosophila* Stock Center | BDSC: 65405 |
| *D. melanogaster*: *Cg-Gal4* (Fat body, Hemocytes) | Bloomington *Drosophila* Stock Center | BDSC: 7011 |
| *D. melanogaster*: *He-Gal4* (Hemocytes) | Bloomington *Drosophila* Stock Center | BDSC: 8699 |
| *D. melanogaster*: *Myo1A-Gal4* (Midgut) | Bloomington *Drosophila* Stock Center | BDSC: 112001 |
| *D. melanogaster*: *phm22-Gal4* (Prothoracic gland) | Gift from Michael B. O'Connor (University of Minnesota) | N/A |
| *D. melanogaster*: *Eip71CD-Gal4* (Epidermis) | Bloomington *Drosophila* Stock Center | BDSC: 6871 |
| *D. melanogaster*: *Lsp2-Gal4* (Fat body) | Gift from Thomas Neufeld (University of Minnesota) | N/A |
| *D. melanogaster*: *tubP-Gal80ts* | Bloomington *Drosophila* Stock Center | BDSC: 7017 |
| *D. melanogaster*: *tubP-Gal80ts* | Bloomington *Drosophila* Stock Center | BDSC: 7019 |
| *D. melanogaster*: *UAS-mCD8::GFP* x 2 | Bloomington *Drosophila* Stock Center | BDSC: 5130, 60698 |
| *D. melanogaster*: *UAS-PGR* RNAi #1 | Bloomington *Drosophila* Stock Center | BDSC: 34035 |
| *D. melanogaster*: *UAS-PGR* RNAi #2 | Vienna *Drosophila* Resource Center | VDRC: v106421 |
| *D. melanogaster*: *UAS-cPges/p23* RNAi | Vienna *Drosophila* Resource Center | VDRC: v106395 |
| *D. melanogaster: UAS-PGR* | This paper | N/A |
| *D. melanogaster*: *UAS-cPges/p23* | This paper | N/A |
| *D. melanogaster*: *UAS-HsPTGES3* | This paper | N/A |
| *D. melanogaster*: *Df(cPges/p23)* | Bloomington *Drosophila* Stock Center | BDSC: 24972 |
| *D. melanogaster: $PGR^{A1}$* | This paper | N/A |
| *D. melanogaster: $PGR^{B1}$* | This paper | N/A |
| *D. melanogaster: $GstS1^{-}$* | This paper | N/A |
| *D. melanogaster: $Mgstl^{-}$* | This paper | N/A |
| *D. melanogaster: $CG33177^{-}$, $CG33178^{-}$* | This paper | N/A |
| *D. melanogaster: $Su(P)^{-}$* | This paper | N/A |
| *D. melanogaster*: *$cPges/p23^{-}$* | This paper | N/A |
| *D. melanogaster*: *SrpHemo-3xmCherry* | Bloomington *Drosophila* Stock Center | BDSC: 78358 |

**S12 Table: Plasmid constructs.**

| Recombinant DNA | Source | Identifier |
|---|---|---|
| cDNA GH27361 | Drosophila Genomics Resource Center | DGRC: 2356 |
| *pBRAcPA-PGR* | This paper | N/A |
| *pUAST-PGR* (UFO02753) | Drosophila Genomics Resource Center | DGRC: 1617200 |
| *pCMV6-HsEP2* | ORIGENE | RC210883 |
| *pBRAcPA-HsEP2* | This paper | N/A |
| *pLV-CMV-aequorin* | VectorBuilder | VB900129-8482vja |
| *pBRAcPA-aequorin* | This paper | N/A |
| pCMV6-GNA15 | Genomics-online | ABIN4214925 |
| *pBRAcPA-Gα15* | This paper | N/A |
| *pBFv-U6.2* | National Institute of Genetics | Kondo and Ueda (2013) |
| *pBFv-U6.2B* | National Institute of Genetics | Kondo and Ueda (2013) |
| cDNA LD23532 | Drosophila Genomics Resource Center | DGRC: 7412 |
| *pCMV6-HsPTGES3* | ORIGENE | RC201254 |
| *pUAST-HsPTGES3* | This paper | N/A |
| *pBRAcPA-HsPTGES3* | This paper | N/A |
| *pBRAcPA-cPges/p23* | This paper | N/A |
| *pUAST-cPges/p23* (UFO02035) | Drosophila Genomics Resource Center | DGRC: 1616694 |
| *pCMV6-HsCOX1* | ORIGENE | RC206436 |
| *pBRAcPA-HsCOX1* | This paper | N/A |
| *pCMV6-HsCOX2* | ORIGENE | RC202245 |
| *pBRAcPA-HsCOX2* | This paper | N/A |

**S13 Table: Primers and oligonucleotides.**

| **Primers for qRT-PCR** | | | |
|---|---|---|---|
| Gene Name | CG Number | Forward (5'-3') | Reverse (5'-3') |
| *Ldh* | CG10160 | CTGGTAGAGTACAGTCCCGA | GGACGAGTCCAAGTTGGTG |
| *Hph* | CG44015 | CGAACAATCTGGCAGCTCAA | GTTCCCGTGGACATGCTAAC |
| *cPges/p23* | CG16817 | GCGAGGGTGATAAGGAGAAGA | TGTTGTGATCGTGTGCTGAC |
| *bnl* | CG4608 | GCCATAACAAGCACACCACA | CTCCTCCAGCTGATCGGAAT |
| *rp49* | CG7939 | AGCTGTCGCACAAATGGCGCAAGC | TTGAATCCGGTGGGCAGCATGTGG |
| **Oligonucleotides for generating *PGR*-targeting gRNAs** | | | |
| Allele | Target ID | Forward (5'-3') | Reverse (5'-3') |
| $PGR^{A1}$ | 2 | CTTCGTAAGTGCAGCTATGCATAG | AAACCTATGCATAGCTGCACTTAC |
| | 3 | CTTCGTCGCTCCAAGTCCATCAAC | AAACGTTGATGGACTTGGAGCGAC |
| $PGR^{B1}$ | 1 | CTTCGCCGCACAACTCAACTACTG | AAACCAGTAGTTGAGTTGTGCGGC |
| | 4 | CTTCGCGCTTAATGCATCCCAGCA | AAACTGCTGGGATGCATTAAGCGC |
| **Primers for screening PGR CRISPR mutants** | | | |
| Allele | | Forward (5'-3') | Reverse (5'-3') |
| $PGR^{A1}$ | | CATTGCTATTGGTCGGCGTTC | CGGTATGAAGATCGAGTGGA |
| $PGR^{B1}$ | | CACCTGAATCACCTGCAATCG | CGGTATGAAGATCGAGTGGA |
| **Primers for generating dsRNAs** | | | |
| *cPges/p23*-1 | Forward (5'-3') | CCGGATCCTAATACGACTCACTATAGCCAAGTCAGGCTGTCAGTCA | |
| | Reverse (5'-3') | CCGGATCCTAATACGACTCACTATAGCACCAGGGCTGTTAAGGAAA | |
| *cPges/p23*-2 | Forward (5'-3') | CCGGATCCTAATACGACTCACTATAGATCATCGATGTCGAATGCAA | |
| | Reverse (5'-3') | CCGGATCCTAATACGACTCACTATAGTAGGCAGCTGGCTTCTTCTC | |
| *EGFP* | Forward (5'-3') | TAATACGACTCACTATAGGATGGTGAGCAAGGGCGAGGAGCCTGTTC | |
| | Reverse (5'-3') | TAATACGACTCACTATAGGCTGGGTGCTCAGGTAGTGGTTGTCGGGC | |
